# Targeted protein evolution in the gut microbiome by diversity-generating retroelements

**DOI:** 10.1101/2024.11.15.621889

**Authors:** Benjamin R. Macadangdang, Yanling Wang, Cora Woodward, Jessica I. Revilla, Bennett M. Shaw, Kayvan Sasaninia, Sara K. Makanani, Chiara Berruto, Umesh Ahuja, Jeff F. Miller

**Affiliations:** Division of Neonatology and Developmental Biology, Department of Pediatrics, David Geffen School of Medicine at the University of California, Los Angeles, Los Angeles, CA, United States; California NanoSystems Institute, Los Angeles, CA, United States; Department of Microbiology, Immunology and Molecular Genetics, University of California, Los Angeles, California, United States; Department of Molecular, Cell, and Developmental Biology, University of California, Los Angeles, Los Angeles, CA, United States; David Geffen School of Medicine, University of California, Los Angeles, Los Angeles, CA, United States; Molecular Biology Institute, University of California, Los Angeles, Los Angeles, CA, United States; Division of Biology and Biological Engineering, California Institute of Technology, Pasadena, CA, United States

## Abstract

Diversity-generating retroelements (DGRs) accelerate evolution by rapidly diversifying variable proteins. The human gastrointestinal microbiota harbors the greatest density of DGRs known in nature, suggesting they play adaptive roles in this environment. We identified >1,100 unique DGRs among human-associated *Bacteroides* species and discovered a subset that diversify adhesive components of Type V pili and related proteins. We show that *Bacteroides* DGRs are horizontally transferred across species, that some are highly active while others are tightly controlled, and that they preferentially alter the functional characteristics of ligand-binding residues on adhesive organelles. Specific variable protein sequences are enriched when *Bacteroides* strains compete with other commensal bacteria in gnotobiotic mice. Analysis of >2,700 DGRs from diverse phyla in mother-infant pairs shows that *Bacteroides* DGRs are preferentially transferred to vaginally delivered infants where they actively diversify. Our observations provide a foundation for understanding the roles of stochastic, targeted genome plasticity in shaping host-associated microbial communities.

## Introduction

Natural selection acts on preexisting genetic variation to favor adaptive phenotypes. In bacteria, this variation primarily arises from mutations and horizontal gene transfer^1^. Nonetheless, genomic integrity is vital for survival and reproductive success, prompting the evolution of mechanisms to channel mutagenesis to localized hotspots within genomes^2–6^ and to limit horizontal transfer^7–9^. Of the known systems that target mutagenesis, diversity-generating retroelements (DGRs) found in bacteria, archaea, and their viruses can generate some of the most extensive repertoires of DNA and protein sequence variants observed in nature^10,11^. Through hypermutation of genes via a mechanism termed mutagenic retrohoming, a single DGR can produce up to 10^30^ unique variable protein sequences^12^. DGR target genes often encode ligand-binding proteins with mutations strategically confined to a discrete subset of codons within a variable repeat sequence (VR) that participates in ligand interactions^13^. The remaining codons in VR, many of which encode structural scaffold residues, remain unmodified^14,15^. This pattern of mutagenesis rapidly diversifies protein function without disrupting structure, leading to readily evolvable ligand-binding capabilities^10^.

In addition to a diversified target gene that encodes a variable protein, a typical DGR includes a template repeat (TR) which is similar but not identical to VR, a uniquely promiscuous reverse transcriptase (RT), and one or more accessory genes (Figure 1A). During mutagenic retrohoming, an RNA intermediate encoded by TR functions as a substrate for reverse transcription by the DGR RT, which selectively mismatches adenine residues. This results in random incorporation of any of the four nucleotides into a cDNA molecule at positions corresponding to TR-adenines. Adenine-mutagenized cDNA is then integrated into VR, replacing the parental allele^16–18^. Because TR is unaltered during this process, and all *cis*- and *trans*-acting factors required for mutagenic retrohoming remain intact, repeated rounds of VR diversification can occur indefinitely to optimize variable protein function.

**Figure 1.**
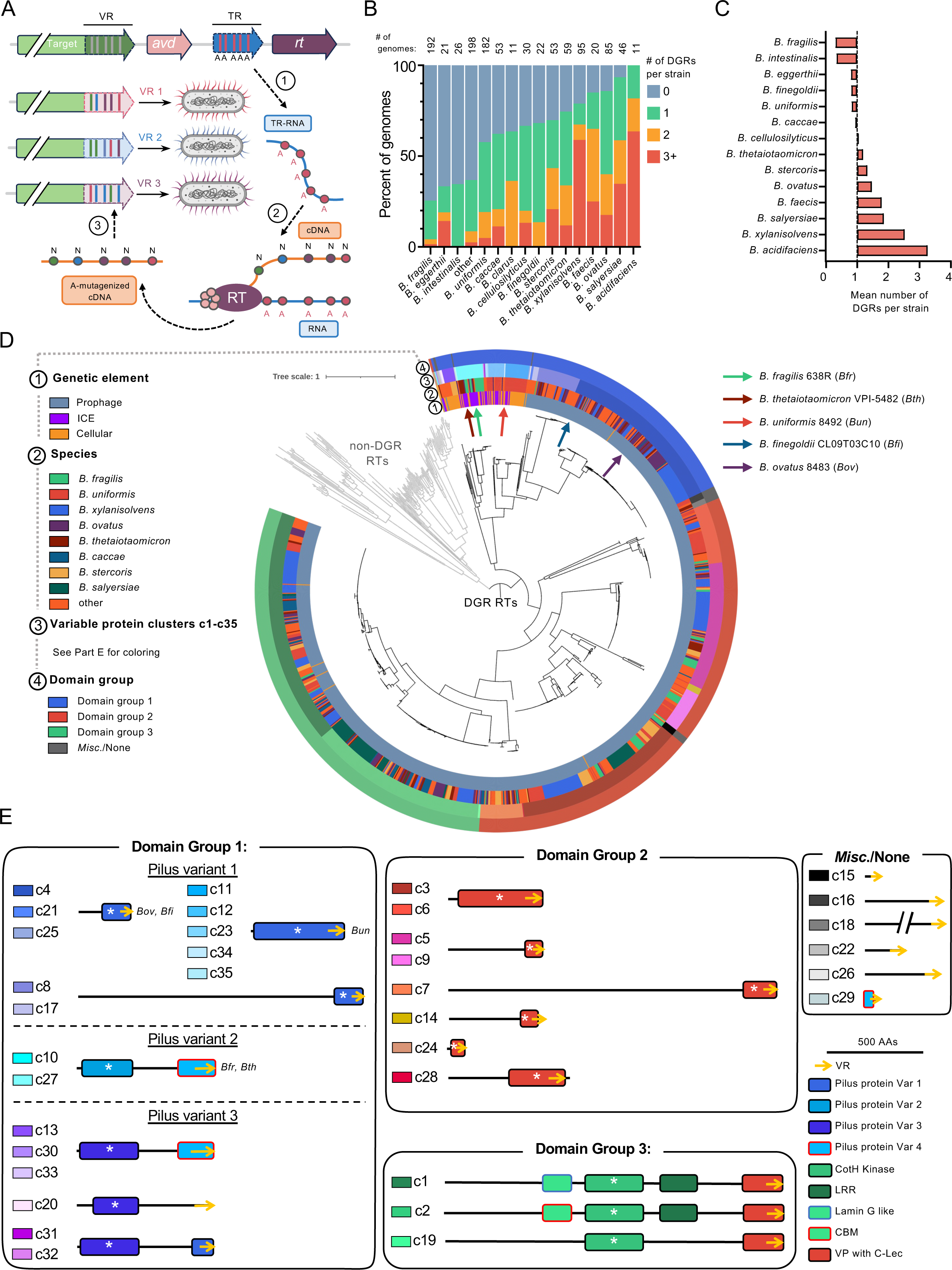
DGRs are widely distributed in *Bacteroides*. (A) Schematic representation of the mechanism of DGR-mediated mutagenic retrohoming. A typical DGR locus is depicted on top. The target gene (green), which encodes a variable protein, contains a VR sequence which is the recipient of mutagenesis. An accessory protein, Avd (pink), helps guide the DGR-encoded RT (purple) to the TR-RNA template. Steps in mutagenic retrohoming include: 1) production of TR-RNA; 2) error-prone cDNA synthesis with misincorporation at TR adenines; and 3) cDNA replacement of parental VR alleles^10,16,17,95^. (B) The number of DGRs found within *Bacteroides* strains, grouped by species. (C) The mean number DGRs per strain grouped by species. The average number of DGRs in all strains (1.01) is set as the baseline. (D) Phylogeny of DGR RTs and non-DGR RTs in *Bacteroides* genomes. Rings depict the genomic location of the DGR (ring 1), species classification (ring 2), variable protein cluster (ring 3), and variable protein domain group (ring 4). Variable protein clusters (c1-c35) and domain groups are colored according to Part E. (E) Visual representation of each cluster, grouped by domain group. The most significant domain is denoted by the white asterix. The color of each cluster corresponds to Part D, ring 3. See also Supplemental Figure S1 and Supplemental Table S4.

The human gastrointestinal (GI) microbiome is enriched in DGRs to an extent that exceeds any other ecosystem characterized to date^19^, yet the dynamics and roles of accelerated protein evolution in this environment have never been systematically interrogated^20^. Strains belonging to the Bacillota phylum and the Fibrobacteres, Chlorobi, and Bacteroidota (FCB) superphylum harbor the vast majority of human microbiome-associated DGRs. As constituents of the FCB group, *Bacteroides* spp. are prominent members of the GI microbiome, forming long-term associations with their human host^21^. They impart health benefits such as secretion of anti-inflammatory molecules^22,23^ and short chain fatty acids (SCFAs)^24^, but are also known to include pathobionts^25^. Their amenability to *in vitro* cultivation and genetic manipulation positions them as a clinically pertinent model for exploring DGR dynamics and function in the context of their natural environment.

Single-nucleotide substitutions such as those produced by DGRs, as well as larger genomic variants such as indels^26^ or the presence of mobile genetic elements (MGEs)^27,28^, can confer characteristics that have profound effects on bacterial phenotypes and fitness. Conventional metagenomic methods commonly lose variant information during assembly and may encounter challenges when categorizing mobile elements during binning^29–47^. Thus, mechanisms underlying accelerated evolution, such as horizontally transferable, DGR-driven mutagenic retrohoming, often evade detection by conventional pipelines. Understanding the functions and phenotypes of complex microbial communities can best be accomplished in the context of extant genotypic variation. To that end, we explored how rapid, targeted evolution by DGRs contributes to *Bacteroides* diversity in the GI tract to gain a foundational understanding of how genome plasticity molds host-associated microbial communities.

Here, we present a systematic analysis of DGR-mediated accelerated evolution in *Bacteroides* species. By analyzing over 1,100 reference genomes, we found that DGRs are prevalent in *Bacteroides* and that a large class of diversified variable proteins have homology to Type V pilins^48^. We show that *Bacteroides* DGRs can be transferred between strains, providing a mechanism for the horizontal transfer of accelerated evolvability. Some *Bacteroides* DGRs are highly active *in vitro* and *in vivo*, and host-encoded factors can further modulate mutagenic retrohoming. In the presence of competition with non-*Bacteroides* strains *in vivo*, diversified *Bacteroides* VR regions can converge to encode similar protein sequences despite being comprised of unique DNA sequences. Finally, by analyzing metagenomic datasets derived from mother-infant pairs, we show that Bacteroidota DGRs are preferentially passed between mothers and infants, where nearly 75% of transferred DGRs adopt a new, predominating VR haplotype. Our results demonstrate that *Bacteroides* DGRs evolve bacterial proteins in the GI microbiome and are active during periods of community instability.

## RESULTS

### DGRs are widespread and diverse in *Bacteroides* species

We analyzed a set of 1,103 *Bacteroides* reference genomes encompassing 47 species from the NCBI RefSeq database^49^ to determine the distribution of DGRs across taxa, to understand their evolutionary relationships and modes of transmission, and to identify functional motifs in the diversified proteins (Supplemental Figure S1A). From this dataset, we found 1,113 unique DGRs distributed across 618 *Bacteroides* isolates (Figure 1B-D). These represented 29 of the 47 species in our dataset and encompassed 11 of the most abundant species in humans (Supplemental Table 4). A solitary DGR was found in 340 isolates (31%), while an additional 278 isolates (25%) contained multiple DGRs, with some genomes of *B. acidifaciens*, *B. xylanisolvens*, and *B. ovatus* harboring up to five unique elements (Figure 1B, Supplemental Figure S1B). DGRs were enriched in these and other species while *B. fragilis* and *B. intestinalis* maintained fewer elements than average (Figure 1C). These results show that DGRs are prevalent and abundant within *Bacteroides* spp. compared to most other bacterial taxa^19,50^.

To gain insight into the relationships between *Bacteroides* DGRs, we built a phylogenetic tree using DGR RT sequences (Figure 1D). Non-DGR RTs identified within the same genome set were included for comparison^51^. We also inspected adjacent sequences for loci related to prophages^52^, integrative and conjugative elements (ICEs)^53^, or plasmids to identify DGRs that are likely to reside on MGEs^19,20^. In the absence of such evidence, DGRs were classified as “cellular” to indicate their presence in cellular genomes. Insights into the potential adaptive roles were obtained by clustering variable proteins and searching for similarities to protein domains of known function^54^. Clusters were further categorized into larger groups based on shared domains with the greatest homology (Figure 1E). For each DGR, information regarding mobility, species, variable protein clustering, and domain groups was overlayed in concentric rings surrounding a DGR RT phylogenetic tree (Figure 1D). As a result of their unique sequence features^10,12^, DGR RTs formed a monophyletic clade distinct from other classes of RTs, consistent with prior studies^17,19,55^. Among the 1,113 DGRs identified, 1055 (95%) resided within predicted MGEs (Figure 1D, ring 1). Variable protein clustering (Figure 1D, ring 3) and domain group relationships (Figure 1D, ring 4) mirrored the phylogenetic patterns of their cognate DGR RT proteins. In contrast, species designations were discordant (Figure 1D, ring 2), supporting the conclusion that *Bacteroides* DGRs evolve as cohesive units, encoding variable proteins that co-evolve with their diversification machinery and are horizontally transferred between strains and species.

None of the *Bacteroides* variable proteins we identified had previously been annotated, therefore, we used profile-based homology to infer their functions^54,56^. VR sequences were uniformly located at the C-termini of variable proteins and were predicted to adopt variant C-type lectin (C-Lec) folds as observed previously^12,19^. Nearly all variable proteins (1092/1113, 98.1%) contained structural domains with homology to one of five broad groups (Figure 1E). Domain group 1 proteins have binding folds similar to Type V pilins expressed by *Bacteroides*, *Porphyromonas,* and related species (PDB: 4EPS, 4QB7, and 5NF4)^48^. This group contains a mixture of prophage-encoded (201/287, 70%), ICE-encoded (13%), and cellular genome-encoded (17%) DGRs. Variable proteins within domain group 2 showed greatest similarity to a *Thermus aquaticus* prophage-encoded diversified protein (TaqVP, PDB: 5VF4)^57^. The absence of other identifiable domains and the observation that group 2 DGRs are found within prophage genomes suggests that these proteins could function as receptor binding components of phage tail fibers. Domain group 3 contains large (>2,000 amino acids) multi-domain variable proteins that include motifs with homology to the active regions of CotH kinases^58^ (PDB: 5JDA), adjacent to other functional domains such as a carbohydrate-binding motifs and leucine-rich repeats (Figure 1E). Overall, *Bacteroides* DGR variable proteins display considerable modularity, whereby diversified C-terminal ligand binding sequences are connected to motifs that are predicted to mediate pilus localization, association with phage tail fibers, signal transduction, or other functions.

### *Bacteroides* DGRs encode pilus subunits and related variable proteins

Based on their conservation and widespread distribution, we reasoned that *Bacteroides* DGRs provide selective advantages to their hosts by accelerating evolution in the gut environment. To explore this, we focused on a selection of five related yet non-identical DGRs present in *B. fragilis 638R (Bfr), B. thetaiotaomicron VPI-5482 (Bth), B. uniformis 8492 (Bun), B. ovatus 8483 (Bov)*, and *B. finegoldii CL09T03C10 (Bfi)* (Figure 1D arrows). Each of these DGRs diversifies variable proteins that share homology to adhesive pilins located at the tips of Type V pili (domain group 1, Figure 1E). Type V pili are modular, extracellular structures composed of anchor, stalk, and tip pilins (Figure 2A), with numerous genes encoding homologs of each subunit type organized into operons spread throughout *Bacteroides* genomes^48^. For example, *Bfr* contains 93 genes clustered into 22 operons that encode components of Type V pili (Supplemental Figure S2A). While the exact roles of these surface appendages in *Bacteroides* spp. are at an early stage of analysis, homologous pili in *Porphyromonas* spp. facilitate coaggregation with other microbes and promote colonization of the oral cavity^59,60^.

**Figure 2.**
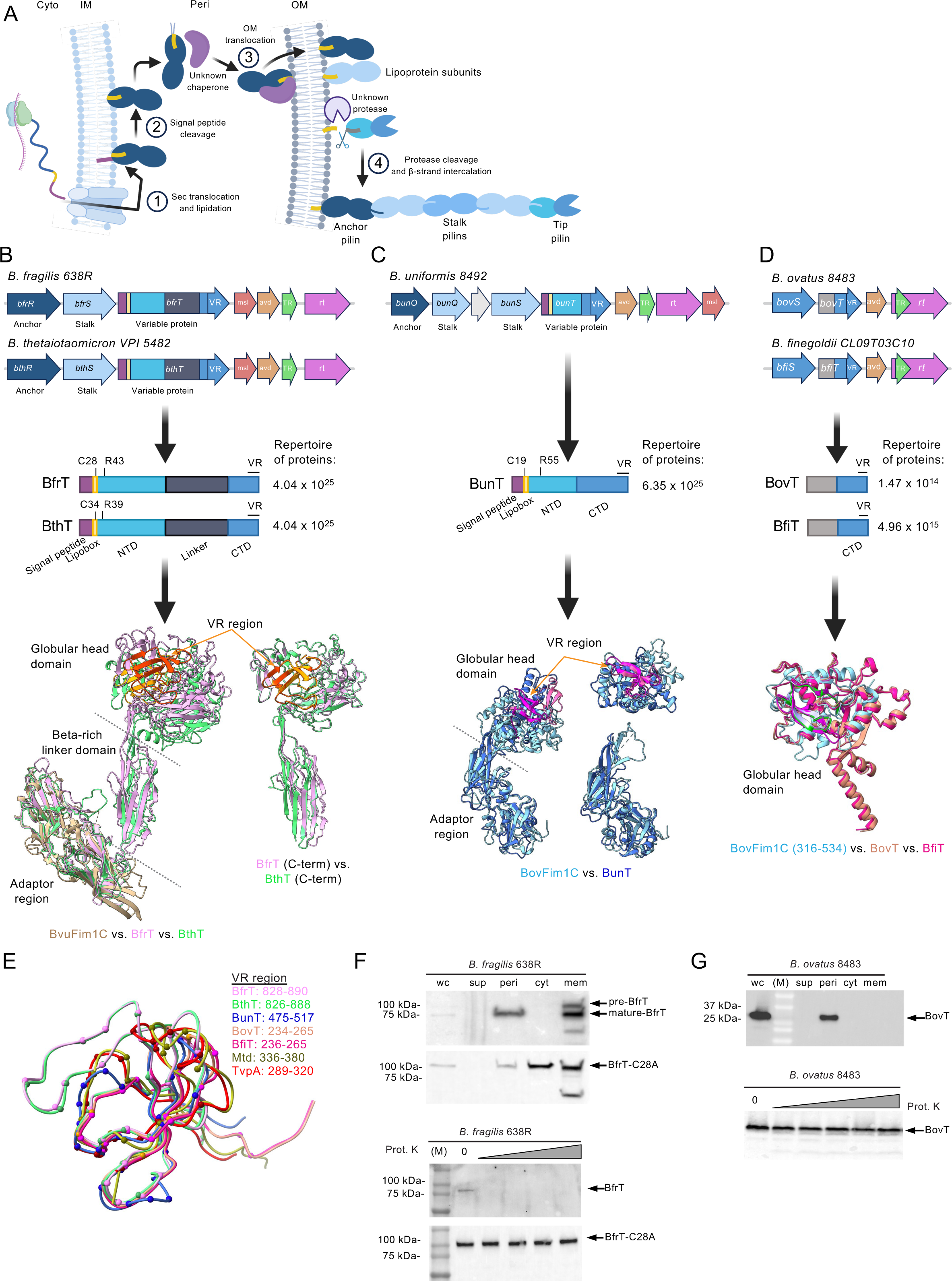
*Bacteroides* DGRs diversify pilus tip adhesins and related proteins. (A) Cartoon representation of the steps in Type V pilus assembly^48^. (1) N-terminal signal peptide recognition and pilus subunit translocation across the inner membrane via the SecYEG translocon, (2) lipidation at a conserved cysteine residue within a lipobox motif and signal peptide cleavage, (3) translocation of the lipidated protein across the outer membrane via an LPP transporter, and (4) incorporation into a growing pilus structure through protease-assisted cleavage which simultaneously releases the protein from the membrane and creates an acceptor site for beta-strand intercalation from an incoming subunit^48,63^. Cyto: cytoplasm; IM: inner membrane; Peri: periplasm; OM: outer membrane; NTD: N-terminal domain; CTD: C-terminal domain. (B-D) DGR loci and their upstream genes in *Bfr*, *Bth, Bun, Bov,* and *Bfi* isolates chosen for this study. Below each loci is a graphical representation of the variable proteins where each domain is colored and labeled with the predicted function. The number of variable proteins that can be generated through mutagenic retrohoming is specified to the right. The protein structures at the bottom are the superposition of the AlphaFold^62^ structures of the variable proteins with known pilus protein structures. BvuFim1C (PDB: 4QB7^48^), BovFim1C (PDB: 4EPS^48^). (E) Superposition of predicted VR-encoded structures for the five *Bacteroides* variable proteins in parts (B)-(D), with the VR-encoded structures of Mtd (gold) and TvpA (red)^13–15^. The positions of variable residues, which are often superimposable in space, are shown as colored balls. (F) Immunoblot of BfrT and BfrT-C28A overexpressed in *Bfr* after cellular fractionation (top) or after intact whole cells were exposed to proteinase K (bottom). See STAR Methods for fractionation protocol and Supplemental Figure S3A for fractionation controls. (G) Immunoblot of BovT overexpressed in *Bov* after cellular fractionation (top) or after whole cells were exposed to proteinase K (bottom). See also Supplemental Figures S2 and S3 and Supplemental Tables S5-8.

The *Bfr, Bth,* and *Bun* variable protein genes (*bfrT*, *bthT*, and *bunT*) are positioned at the ends of operons predicted to encode anchor and stalk proteins, which link tip pilins to the bacterial cell surface (Figure 2A-C, Supplemental Figure S2B). The *Bov* and *Bfi* elements are located adjacent to prophage genes, but it is unclear if they are carried by a phage (Figure 1D). All five *Bacteroides* DGRs encode an Avd-like protein^61^, and *Bfr, Bth*, and *Bun* encode an additional accessory factor, Msl (<u>M</u>ut<u>S</u>-like), with homology to the mismatch recognition domain of MutS^12^ (Figure 2B-D). Each of the five TRs contains 30 to 45 adenines, providing the capacity to generate massively diverse repertoires of 10^18^-10^27^ potential VR DNA sequences, and 10^14^-10^25^ different polypeptides at the C-termini of their cognate variable proteins.

We generated predicted 3D structures with high per-residue confidence scores (pLDDT >90) for all five variable proteins and compared them to known atomic structures of *Bacteroides* pili (Figure 2B-E)^48,62^. BfrT and BthT, which share 61% amino acid identity (AAI) with each other, displayed a hybrid pilin structure with three domains. Their N-terminal domains (NTDs) are homologous to the NTD of BvuFim1C, a structurally characterized Type V stalk pilin encoded by *B. vulgatus*^48^ (Figure 2B, Supplemental Table 5,6). Their C-terminal domains (CTDs), which contain DGR-diversified residues organized in a C-Lec fold, diverge from BvuFim1C and instead adopt a globular head structure similar to ligand-binding CTDs of tip pilins (Figure 2B). Interestingly, both BfrT and BthT exhibited an additional third domain that connects stalk and tip domains together. The *Bun* DGR variable protein, BunT, adopts a canonical bipartite pilin structure, with N- and C-terminal domains that share high homology to BovFim1C (Figure 2C, Supplemental Table 5)^48^, a Type V tip pilin encoded by *B. ovatus*. While the globular head of BovFim1C is static, the BunT globular head displays diversifiable VR-encoded residues (Figure 2C). The *Bov* and *Bfi* variable proteins, BovT and BfiT, are comparatively small, highly similar to each other (86.5% AAI), and fold into a structure that is homologous to the CTD globular head of BovFim1C (Figure 2D), with an N-terminal pair of alpha helices in place of the pilin-like NTD (Figure 2D, Supplemental Table 7). Comparing predicted structures of *Bacteroides* VPs with structurally characterized DGR VPs^14,15^ revealed a remarkable superimposition of the overall tertiary structure, including overlap in the spatial locations of variable residues (Figure 2E) despite substantial differences in amino acid sequences, providing evidence for the conservation of similar ligand-binding interactions^13^.

Biogenesis of a Type V pilus is a multistep process that involves: *1)* pilin translocation and lipidation, *2)* signal peptide cleavage, *3)* translocation to the outer membrane, and *4)* incorporation into a growing pilus^48,63,64^ (Figure 2A). As shown in Figure 2B-C, BfrT, BthT, and BunT encode conserved N-terminal signal sequences, lipobox motifs, and protease-cleavable arginines required for pilus assembly. We placed affinity tags^65^ at the C-termini of BfrT and BfrT-C28A, a mutant derivative lacking the conserved cysteine required for lipidation and translocation to the outer membrane^64^, and expressed the tagged proteins in *Bfr*. Following induction and cell fractionation (Supplemental Figures 3A, B, Supplemental Table 8), BfrT was readily detectable in membrane fractions in both pre-processed and mature forms, and in the periplasm as the mature form (Figure 2F). Treatment of intact cells with proteinase K resulted in digestion of BfrT, consistent with its localization on the cell surface, while the C28A mutant was protease resistant (Figure 2F). Additionally, immunofluorescence demonstrated BfrT on the cell surface that was sensitive to proteinase K, but no cell surface staining of the C28A mutant was observed (Supplemental Figure S3C). Mass spectroscopy of mature BfrT showed that cleavage had occurred at R43 (Supplemental Figure S3D, E), as predicted. We next examined the localization of tagged BovT and BfiT, both of which were found exclusively in the periplasm and were resistant to proteinase K (Figure 2G). These observations identify two classes of variable proteins within our five *Bacteroides* strains. DGRs belonging to *Bov* and *Bfi* diversify periplasmic proteins that are structurally related to tip adhesins, but either require additional factors for incorporation into pilus structures or have evolved to perform different functions dependent on their ligand-binding capabilities, whereas *Bfr, Bth,* and *Bun* DGRs diversify Type V pilus tip adhesins.

### Horizontal transfer of *Bacteroides* DGRs

Horizontal transfer of DGRs that diversify phage tail fiber proteins has been well studied^17,18^. In contrast, the mobility characteristics of DGRs that target bacterial proteins are relatively unexplored. To address this, we exploited the observation that the DGRs encoded by *Bfr* and *Bth* are flanked by mobility and transfer genes characteristic of ICEs (Figure 3A). Like phage, ICEs often confer selective advantages to their hosts by carrying cargo that encode colonization factors, metabolic capabilities, virulence determinants, antibiotic resistance, or other accessory functions^66–73^.

**Figure 3.**
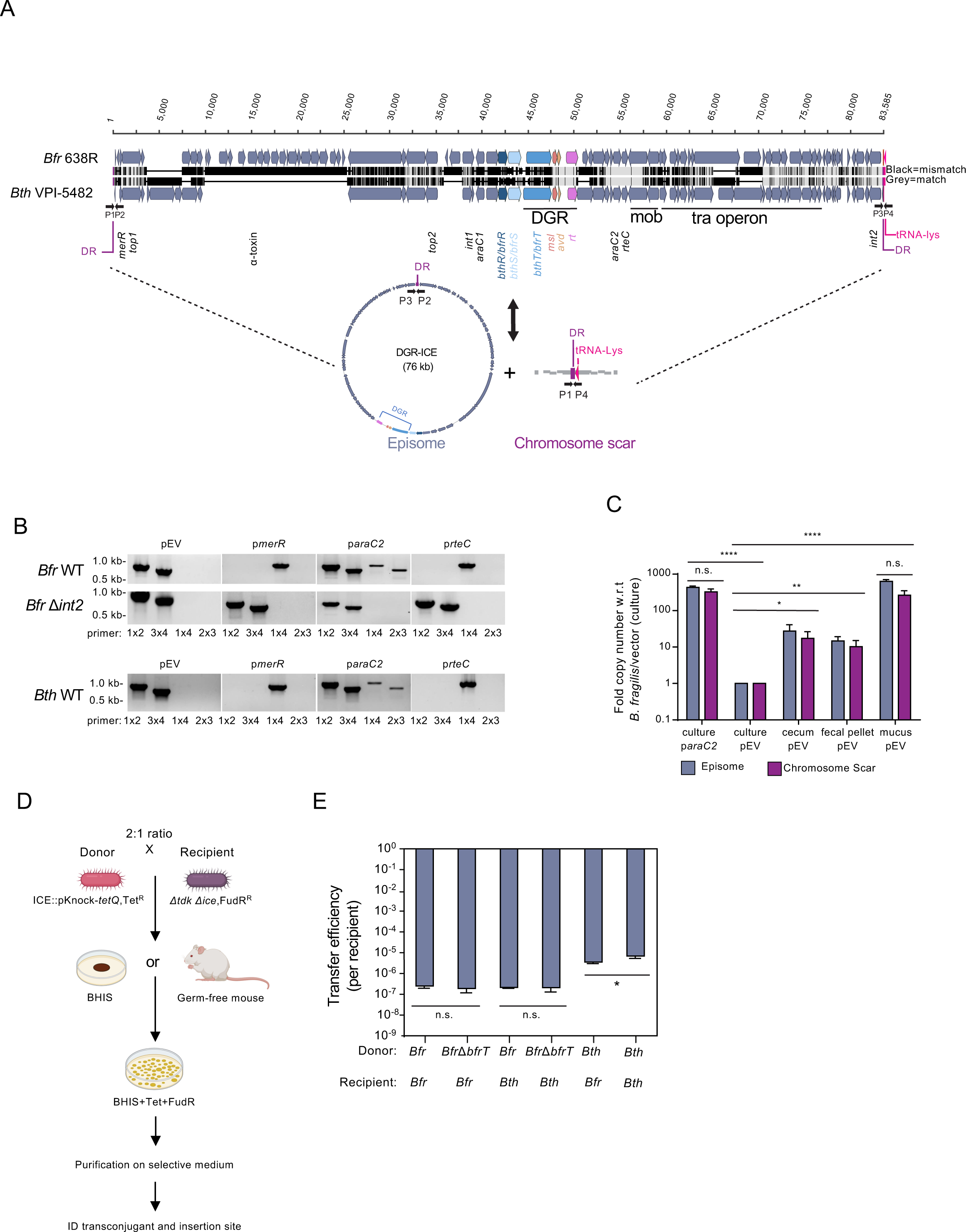
*Bacteroides* DGRs are horizontally transferred between strains and species. (A) (Top) Synteny between *Bfr* and *Bth* ICEs. Important elements within each ICE, including the DGR loci, are labeled. (Bottom) Cartoon schematic of the *Bfr* ICE conversion between chromosomally integrated and episomal forms. Binding of primers P1-P4 shown for integrated ICE, episomal ICE and chromosomal scar. (B) PCR products from *Bfr* (top) and *Bth* (bottom) using primers to differentiate chromosomally integrated ICEs from excision products (episome and scar) following overexpression of designated ICE encoded regulatory genes. pEV, empty vector. (C) Ratios of excised episomes or chromosomal scars to integrated ICEs, normalized to pEV containing strains cultivated *in vitro.* Samples from cecum, fecal pellet, and mucus originate from monocolonized SW mice (n=3). (D) Experimental setup for ICE mating assays. (E) The transfer efficiency, defined as the number of transconjugant cells divided by the total number of recipient cells, of *in vitro* ICE mating assays. Statistical analysis: *p<0.05, **p<0.01, ***p<0.001, ****p<0.0001. ANOVA test with Holm-Sidak multiple comparison correction. Error bars, standard deviation of mean. See also Supplemental Figure S4 and Supplemental Tables S9-13.

We identified 41 DGRs residing within ICEs present in 10 different *Bacteroides* species, including the *Bfr*, *Bth, and Bun* elements in Figure 2B,C, (Supplemental Table 4). The majority of target genes diversified by these DGRs (38/41) encode variable proteins from domain group 1 that are predicted to function as Type V pilus tip adhesins and are encoded directly downstream from anchor and stalk subunits (Supplemental Figure S4A). Of these 41 ICEs, 16 aligned closely with each other and shared conserved features including conjugation and DNA integration genes (*tra, int*), homologs of known transcriptional regulators (*merR*, *rteC, araC* family members)^74^, and direct repeats resulting from site-specific chromosomal integration into a tRNA-Lys locus (Figure 3A, Supplemental Figure S4A). In addition to related variable proteins, the DGRs encoded by these ICEs share similar TRs and RTs (Supplemental Tables 9 and 10), suggesting they disseminated among *Bacteroides* species via horizontal transfer of an ancestral DGR-encoding ICE.

To measure ICE activity, we developed PCR assays to differentiate integrated *vs.* excised forms of the *Bfr* and *Bth* elements (Figure 3A). When WT cells were grown *in vitro*, only the integrated form of the ICEs could be detected (Figure 3B). This was not unexpected, given that mobile genetic elements often require environmental signals to induce their mobility^74^. We next identified ICE-associated regulatory loci (Figure 3A) and created strains that ectopically express each regulatory factor (Supplemental Table 11). Overexpression of *araC2* induced ICE excision and circularization, as both circular episomes and chromosomal scars were observed in *Bfr* and *Bth* (Figure 3B). Overexpression of *merR* or *rteC*, however, resulted in the absence of both integrated forms and episomes, indicating the loss of ICEs from these cells (Figure 3B and Supplemental Table 11). Efficient excision in the absence of integration presumably leads to episome segregation during replication. To further identify requirements for excision and integration, we individually deleted ICE-encoded integrase (*int1, int2*) and topoisomerase (*top1, top2*) genes. Excision of the *Bfr* ICE was dependent on the presence of an intact *int2* integrase gene (Figure 3A, B), but was unaffected by knocking out *int1, top1,* or *top2* (Supplemental Table 12). Finally, to determine if inducing signals are provided *in vivo*, we colonized germ-free Swiss Webster mice with *Bfr* and measured relative levels of episome and chromosomal scar formation by qPCR in bacteria recovered from fecal pellets, cecal contents, and colonic mucosa. As shown in Figure 3C, compared to *Bfr* plus empty vector cultured *in vitro*, we observed 17- and 27-fold increases in ICE activity in fecal pellets and cecal content, respectively, and a nearly 700-fold increase in colonic mucus, the natural habitat of *Bacteroides*.

Resident ICEs are known to exclude integration by homologous mobile elements^75,76^. To create recipient cells suitable for mating experiments, we overexpressed *merR* to promote ICE excision and loss (Figure 3B), curing *Bfr* and *Bth* strains of their DGR-containing ICEs (ΔICE) and leaving their chromosomal integration sites free. In matings between isogenic WT donor and ΔICE recipients, transconjugants were isolated at a frequency of 10^-5^ to 10^-7^, with *Bth* donors displaying greater transfer efficiencies than *Bfr* donors (Figure 3D, E, Supplemental Table 13). To determine if transfer requires the DGR-encoded variable protein, we deleted *bfrT* and found that *Bfr*Δ*bfrT* mutants displayed the same transfer efficiencies as the WT parent, demonstrating that the diversified Type V pilus tip adhesin is dispensable for horizontal transfer. ICE conjugation was also observed in gnotobiotic mice (Supplemental Figure S4B,C). Our results demonstrate that DGRs in *Bfr* and *Bth* are encoded within functional ICEs that undergo conjugative transfer *in vitro* and *in vivo*, providing an explanation for the phylogenetic distribution of DGRs in *Bacteroides* and a mechanism for the horizontal transfer of accelerated protein evolution between species.

### Differential control and mechanistic conservation of mutagenic retrohoming

To characterize the real-time dynamics and mutational patterns of *Bacteroides* DGRs, we measured mutagenic retrohoming levels *in vitro* and *in vivo* and interrogated the diversified sequences. Strains carrying the DGRs shown in Figure 2B-D were grown *in vitro*, sampled over a two week period, and VRs were barcoded, amplified, and deep sequenced (Figure 4A). We calculated the percentage of VRs that had diverged from their parental sequence and found that the *Bov* and *Bfi* elements showed remarkably high levels of mutagenesis, with 13% or 40% of VRs diversified by day 14, respectively (Figure 4B). This was unexpected, since the activity of DGRs that mutagenize bacterial genes in other genera has been reported to be low or absent during *in vitro* growth, reflecting their apparent regulation^10,19^. VR mutagenesis was abolished in strains harboring knockout mutations in *rt* (Δ*rt*) (Figure 4C, Supplemental Figure S5A) and restored by complementation with wild type *rt* expressed at an ectopic location (Supplemental Figure S5B). In contrast, VR sequences from *Bfr*, *Bth,* and *Bun* displayed *in vitro* levels of mutagenesis that were 100- to 10,000-fold lower than *Bov* or *Bfi* (Figure 4B). To measure DGR activity *in vivo*, we monocolonized germ-free Swiss Webster mice with individual *Bacteroides* strains. Colonization levels in the GI tract were similar for each *Bacteroides* strain as measured by colony-forming units in fecal pellets (Supplemental Figure S5C). *Bov* and *Bfi* DGRs were highly active in the murine GI tract, displaying levels of diversity similar to those observed *in vitro*, while the activity levels observed with *Bfr*, *Bth*, and *Bun* remained low (Figure 4B). Next, we used RNA-Seq to probe relationships between mutagenic retrohoming and transcription of DGR-encoded genes. We observed significantly higher relative amounts of transcripts encoding *avd,* TR, and *rt* in high activity strains (*Bov* and *Bfi*, Figure 4D) compared to those with low DGR activity (*Bfr, Bth, Bun*), suggesting that DGR mutagenesis is regulated, at least in part, at the transcriptional level. Thus, mutagenic retrohoming levels in our five *Bacteroides* strains fall into two categories. DGRs carried by *Bov* and *Bfi* are constitutively active, while those in *Bfr, Bth,* and *Bun* appear to be tightly regulated, as commonly observed in other systems^19,77^.

**Figure 4.**
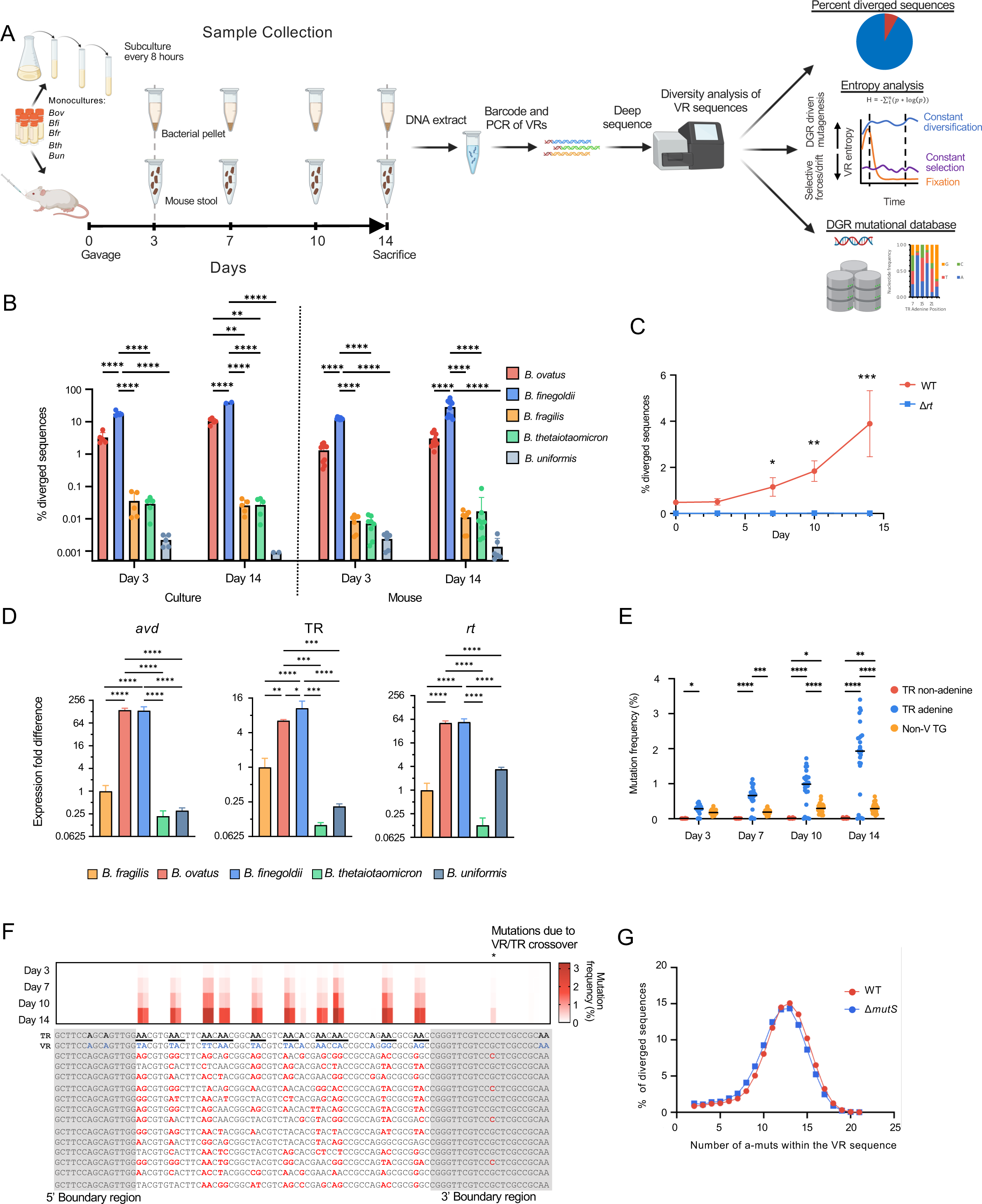
*Bacteroides* DGRs are differentially active *in vitro* and *in vivo*. (A) Schematic of the experimental design. (B) Percent of VRs that mutated from the parental VR sequence in cells grown *in vitro* or present in stool samples from monocolonized SW mice (n=5-11). Error bars, standard deviation of mean. (C) Percent of VR sequences that diverged from their cognate parental VRs in *Bov* WT and *Bov* Δ*rt* in stool samples from monocolonized SW mice (n=4). Error bars, standard deviation of mean. (D) Expression of DGR encoded genes *avd*, TR, and *rt* measured by RNA-Seq and normalized to *gyrA,* from mid-log phase cells grown *in vitro* (n=4). Error bars, standard deviation of mean fold change. (E) Mutation frequencies of individual nucleotide positions within the *Bov* target gene over a two week period from a population of VRs extracted from stool samples of monocolonized mice (n=4). (F) Nucleotide entropy at each position along the *Bov* VR for a population of VRs extracted from stool samples of monocolonized mice (n=4). Underneath the graph are the parental TR and VR sequences followed by representative VR sequences at Day 14. TR adenines are bolded, positions in VR corresponding to TR adenines are in blue, and positions in the mutated VR sequence that are different from the parental VR are shown in red. The gray boxes show the presumed 5’ and 3’ boundary regions where no adenine mutations are observed. (G) Distribution of the number of nucleotide substitutions per VR read in *Bov* WT *vs. Bov ΔmutS* cells grown *in vitro*. Statistical analysis: *p<0.05, **p<0.01, ***p<0.001, ****p<0.0001. ANOVA test with Tukey multiple comparison correction (B,D,G) or Holm-Sidak multiple comparison correction (C). See also Supplemental Figures S5.

Mutagenic retrohoming in *Bacteroides* demonstrated remarkable specificity for substitutions at TR adenines (Figure 4E), a hallmark of DGR RT enzymes observed across taxa^16,18,77^. To identify positional effects, we exploited the constitutive activity of the *Bov* and *Bfi* DGRs to examine time-dependent levels of mutagenesis as a function of position within VR (Figure 4F, Supplemental Figure S5D). The central region of VR is enriched with sequences corresponding to TR AAC motifs (Figure 4F, underlines) that enable random substitution at one or both adenines^12,20^, accounting for the significant accumulation of mutations over time. In contrast, VR positions at the 5’ and 3’ ends displayed minimal to no mutations despite the presence of TR adenines, consistent with boundary effects similar to those reported for the *Bordetella* phage BPP-1 DGR^16^, including a crossover interval near the center of the 3’ boundary region where mutagenized cDNA integrates to replace parental VRs. Finally, by counting the number of adenine mutations in uniquely diversified VR sequences, we observed a median mutational density of about 50% of available positions (Figure 4G, Supplemental Figure S5F). A nearly identical mutational density was reported in *Bordetella in vivo*^78^, and *in vitro* with cDNA synthesized by purified BPP-1 RT, Avd, and TR-RNA in the presence of dNTPs^79,80^. These observations highlight the striking conservation of the mechanisms of adenine-specific mutagenesis and cDNA integration in distantly related DGRs and bacterial hosts.

### *Bacteroides* DGRs preferentially create non-synonymous substitutions that alter side chain chemistry

The high levels of mutagenic retrohoming conferred by the *Bov* and *Bfi* DGRs allowed us to build the largest dataset of experimentally derived diversified VR sequences available to date. VR-encoded amino acids that differed from their initial parental sequence during serial subculturing were classified as synonymous or non-synonymous, and the amino acid substitution frequency at each codon position was calculated. Non-synonymous substitutions at cognate TR adenine positions predominated our dataset, accounting for 99.9% of amino acid changes resulting from nearly 4,500,000 DGR-generated VR mutations (Figure 5A, Supplemental Figure S6A-C). The vastly disproportionate number of non-synonymous substitutions generated by DGR mutagenesis arises directly from the bacterial genetic code, coupled with a high abundance of TR AAY (AAC or AAT)^15^ motifs, which account for 10 of the 12 variable codons between the 5’ and 3’ boundaries of the *Bov* VR (Figure 4F). The abundance of TR AAY motifs extends beyond *Bov.* In our *Bacteroides* dataset, AAY motifs (∼10.5/TR) outnumber single-adenine motifs (∼5.1/TR) (Supplemental Figure S6D,E Mann-Whitney p<0.0001), underscoring their significance. For AAC motifs, 16 potential codons can be generated through random adenine mutagenesis, 14 of which encode unique amino acids (Figure 5B), while only two codons are synonymous (AGC and TCC which encode serine), and a stop codon can never be generated. The side chains of the 15 amino acids produced by AAC mutagenesis encompass the entire range of available chemical properties, including polar uncharged, small hydrophobic, large hydrophobic, positive charge, negative charge, and no side chain (glycine) (Figure 5B). Therefore, the predominance of TR AAY motifs leads to an expansive and chemically diverse list of amino acids at variable sites and an overwhelmingly high probability of producing non-synonymous mutations due to adenine mutagenesis.

**Figure 5.**
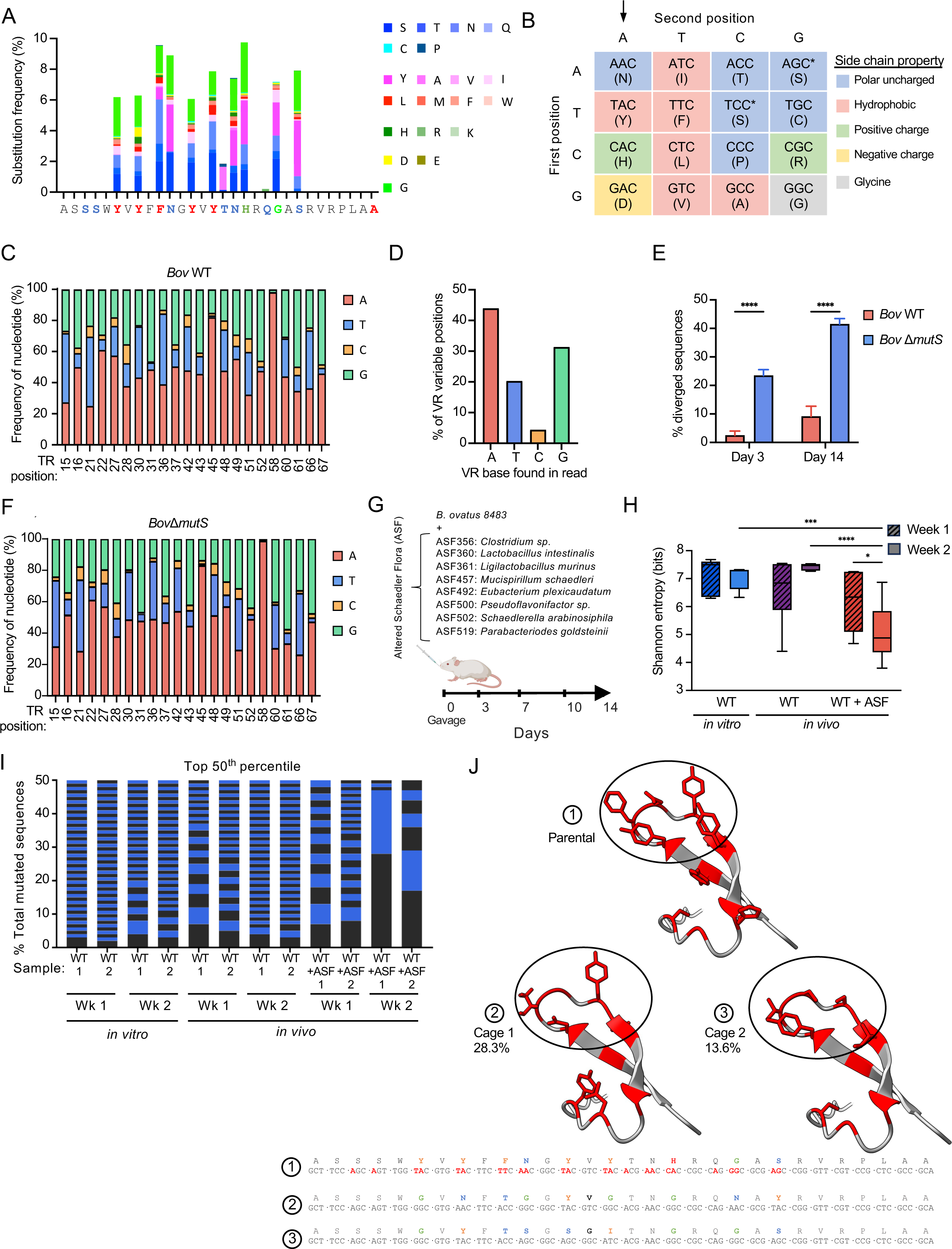
*Bacteroides* DGRs are poised to alter amino acid side chain chemistry and respond to competition *in vivo*. (A) Substitution frequency at *Bov* VR codons that correspond to TR adenines from cells grown *in vitro*. (B) Table showing codons that can be generated through mutagenic retrohoming of a TR AAC motif, colored by the chemical class of the amino acid side chain. (C) Nucleotide frequency at individual variable VR positions within diversified *Bov* VRs. The corresponding TR adenine position is listed below each bar. (D) Cumulative frequency of each of the four nucleotides at variable VR sites within diversified *Bov* VRs. (E) Percent of VRs that mutated from the parental VR sequence in *Bov* WT and *Bov* Δ*mutS* cells grown *in vitro*. (F) Nucleotide frequency at individual variable VR positions within diversified *Bov* Δ*mutS* VRs. The corresponding TR adenine position is listed below each bar. (G) Co-colonization of germ free mice with *Bov* with or without Altered Schaedler Flora (ASF), an 8 member bacterial community. (H) Shannon entropy of diversified VR sequences obtained from *Bov* cells grown *in vitro* or present in fecal samples of gnotobiotic mice (n=5-11). *p<0.05, **p<0.01, ***p<0.001, ANOVA with Tukey multiple comparison correction. (I) Graphical representation of diversified *Bov* VR populations derived from cells grown *in vitro* or present within fecal samples of gnotobiotic mice. The height of the horizontal bars represents the frequency of appearance, with alternating black and blue indicating unique VR sequences. (J) VR encoded predicted structures of the parental *Bov* VR and two of the most commonly observed VR sequences derived from fecal samples of separately caged gnotobiotic mice colonized with *Bov* plus ASF bacteria. The VR nucleotide and amino acid sequences are shown below and the frequency of the individual VR sequence within the diversified VR population is indicated. Also see Supplemental Figure S6.

On closer examination, we noticed a non-random pattern of diversified residues whereby the chemical property of the side chain often switched during mutagenesis (Supplemental Figure S6F). By positioning an adenine in the middle of the codon, random mutagenesis of the first position results in four potential codons, each with unique side chain chemical properties (Figure 5B, arrow). Interestingly, we observed a non-random pattern in the nucleotide frequency at variable positions. Adenine was the most prevalent nucleotide at the majority of the variable positions in both *Bov* (Figure 5C, D) and *Bfi* (Supplemental Figure S6G, H), followed by guanine and thymine, while cytosine was rarely observed. When specifically focusing on VR positions corresponding to TR AAC motifs, an adenine was observed in the second position at very high frequencies that were often greater than 50%. Thus, the nucleotide frequency across variable positions in VR exhibits stochastic but non-uniform patterns, with a clear bias to incorporate adenines that result in frequent switching of amino acid side chain chemistry.

Mechanistic models of mutagenic retrohoming predict the formation of heteroduplexes between mutagenized cDNAs and parental VR sequences with an unusually high density of mismatches^10,78,80^. Taking advantage of the constitutive activity of the *Bov* DGR, we explored the impact of mismatch repair on the level and pattern of VR mutagenesis by knocking out *mutS*. *Bov*Δ*mutS* exhibited substantially higher levels of VR mutagenesis than WT, with up to 25% and 43% of VRs mutated by days 3 and 14, respectively, corresponding to an approximate 5- to 10-fold increase compared to the parent strain (Figure 5E). Complementation with intact *mutS* restored mutagenic retrohoming to WT levels (Supplemental Figure S6I). Intriguingly, analysis of nucleotide frequencies at mutated variable sites revealed an almost identical distribution (Figure 4G) and pattern (Figure 5C, F) when diversified VRs were compared between *Bov* WT and *Bov*Δ*mutS*. These results support a mechanism in which MutS-mediated repair operates in an all-or-nothing manner, whereby VR heteroduplexes are either converted back to the parental sequence or fully escape mismatch repair.

### DGR dynamics under competitive pressure

Estimating the percentage of VRs that have undergone mutagenic retrohoming provides an incomplete picture of the true extent of sequence diversity. For example, a population in which a majority of diverged VRs encode the same or a limited number of DNA or protein sequences would clearly be distinct from one in which most of the diverged sequences differed from each other. Thus, we calculated the Shannon entropy^81^ of VRs that had undergone adenine-mutagenesis under different conditions. Although this metric can be biased at small sample sizes^82^, our large mutational dataset allows it to encompass the relative strengths of two opposing forces: i) mutagenic retrohoming (Supplemental Figure S6J), which increases VR entropy by randomizing sequences, and ii) purifying selection, which decreases entropy through preferential propagation of mutagenized VR sequences that provide a competitive advantage^19^.

We measured VR entropy in populations of *Bov* that were grown *in vitro* or in germ-free mice colonized with or without Altered Schaedler Flora (ASF), an eight-member bacterial consortium, to provide interspecies competition^83^ (Figure 5G). As expected, VR sequences derived from *in vitro* grown cells displayed high levels of amino acid entropy at early and late timepoints (Figure 5H), indicating that mutagenic retrohoming was primarily driving VR diversity. When *Bov* was introduced into germ-free mice, VR entropy displayed high values similar to *in vitro* samples, suggesting the absence of strong selective forces in monocolonized hosts. In contrast, VR sequences from animals co-colonized with both Bov and ASF displayed a time-dependent decrease in entropy that became highly significant by week 2 post-gavage, indicative of positive selection. On closer examination, samples from *in vitro* grown or monoassociated *Bov* contained VRs that were almost entirely different from each other, while samples from mice co-colonized with ASF contained populations in which two to three unique VRs comprised >50% of all mutated sequences (Figure 5I). Furthermore, in separately caged co-colonized mice, mutated populations of VRs had converged by day 14 to express similar amino acid sequences despite being encoded by different DNA sequences. In the examples shown in Figure 5J, several hydrophobic side chains have been replaced by polar, uncharged residues while maintaining a hydrophobic interaction site near the top of the VR, creating a more open binding pocket and suggesting these variant sequences have evolved to interact with a new, common ligand. Taken together, these observations demonstrate that in the face of competition with other microbes, VR entropy decreases in a manner expected for environmental conditions that exert positive selection.

### DGRs are active during the intergenerational handoff from mothers to infants

We hypothesized that dynamic changes that accompany the intergenerational handoff of gastrointestinal microbes from mothers to infants during birth could select for DGR-mediated adaptations that facilitate colonization and persistence in a new host. To explore the role of DGRs during this period, we performed an integrative analysis of retroelements present in metagenomic datasets of human fecal microbiomes from 144 longitudinally sampled mother-infant pairs^84–86^ as well as from a dataset of 146 healthy adults^87^ (Supplemental Figure S7A). We identified 5106 different DGRs, of which 2740 were identified within mother-infant pairs, with 698 DGRs found uniquely in infants, 1654 found only in mothers, and 388 DGRs that were apparently transmitted from mothers to infants during or after birth (Figure 6A, Supplemental Table S14). To gain a deeper understanding of these elements, we analyzed DGR phylogeny, variable protein domains, and predicted transfer vector, similar to our prior analysis with *Bacteroides* (Figure 1) but expanded to include all phyla (Figure 6B).

**Figure 6.**
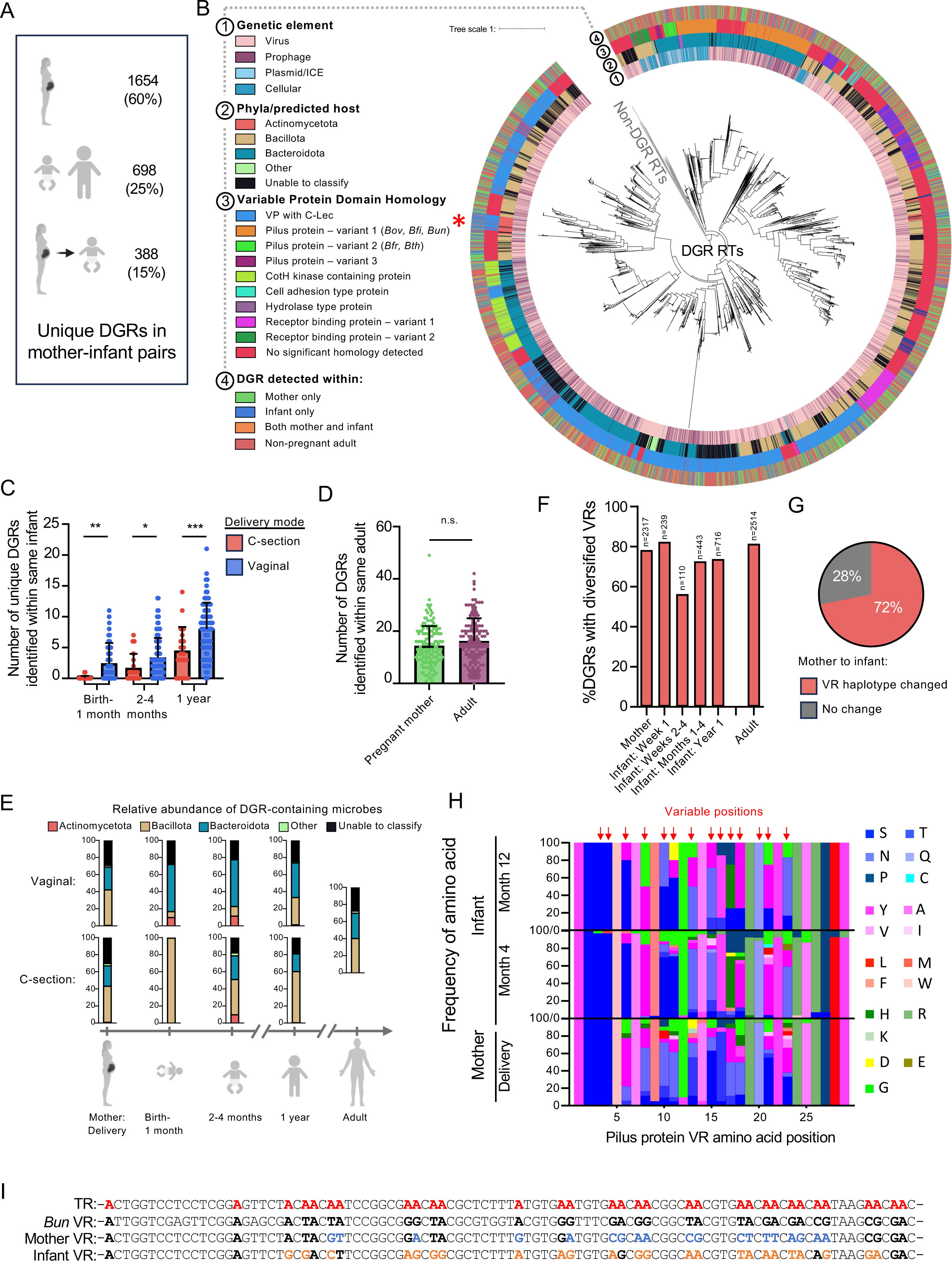
DGRs undergo a burst of activity when transmitted from mother to infant. (A) Number of unique DGRs found in mothers, in infants, or transmitted from mother to infant. (B) Phylogeny of DGR RTs found in metagenomes derived from mothers, infants, and healthy adults^84,85,96,97^. Rings depict the genomic location of the DGR (ring 1), predicted host phyla (ring 2), variable protein homology (ring 3), and classification of the DGR host (ring 4). Red asterix shows an area with Actinomycetota-harboring DGRs found primarily in infants. (C) Number of DGRs identified per infant, grouped by age and mode of delivery. *p<0.05, **p<0.01, ***p<0.001, ANOVA with Holm-Sidak multiple comparison correction. (D) Number of DGRs identified per adult. n.s.: not significant. (E) Taxonomic distribution of DGR-containing microbes, grouped by age and mode of delivery. (F) Percent of DGRs with mutations at VR positions that correspond to TR adenines, grouped by age. (G) Percent of DGRs where the predominant VR haplotype sequence changed between mother and infant (n=388). (H) Frequency of VR encoded amino acids from a DGR that was transmitted from mother to infant. The target gene of this DGR is predicted to encode a Type V pilus tip adhesin. (I) TR and VR sequences of *B. uniformis* 8492 (*Bun*) and the VRs from a similar DGR identified within a mother and her infant. Red, *Bun* TR adenines; Bolded, *Bun* VR nucleotides that correspond to TR adenines; Blue, VR nucleotides that differ between *Bun* and the maternal VR sequence; Orange, VR nucleotides that differ between the maternal and infant VR sequences. See also Supplemental Figures S7 and Supplemental Tables S14-15.

The majority of DGRs in this dataset were identified as phage- or prophage-encoded (93%). Of the 344 DGRs found in cellular genomes, plasmids, or ICEs, 214 (62%) were predicted to reside within Bacteroidota, while 106 (31%) were classified as Bacillota. Cellular or ICE-encoded Bacteroidota DGRs almost exclusively diversify pilus subunits or other cell adhesion proteins, as observed with the dataset analyzed in Figure 1B. In contrast, most Bacillota variable proteins classified as cellular, plasmid, or ICE-encoded have domains similar to phage receptor binding proteins, suggesting this binding module was co-opted to perform some other function. DGRs in this dataset were distributed throughout infants, mothers, and adults, except for a group of DGRs belonging to Actinomycetota (Figure 6B, red star), which were enriched in infants, and to a lesser extent in mothers, but rarely found in nonpregnant adults.

Mode of delivery had a significant impact on the number of DGRs identified in infants (Figure 6C). At birth, DGRs were much more common in vaginally born infants compared to those born by C-section, and this trend persisted throughout the first year of life. The mean number of unique DGRs in infants at one year was significantly less than the mean number of DGRs in mothers and adults (Figure 6C,D, p<0.0001 all comparisons of 1-year-olds vs adults), showing that new DGRs continue to be acquired throughout life. Mode of delivery also correlated with the relative distribution of DGR-containing taxa. Prior to delivery, there were no significant differences in the taxonomy of microbes harboring DGRs in mothers undergoing C-section compared to mothers delivering vaginally (Figure 6E). However, infants delivered vaginally showed a much higher proportion of DGR-containing Bacteroidota, while infants born via C-section acquired an initial set of DGRs that were almost exclusively encoded by Bacillota. By the end of infants’ first year, the taxonomy of DGR-containing microbes more closely resembles the distribution found in adults regardless of the mode of delivery (Figure 6E). Comparisons between breastfed and formula fed infants or between males and females revealed no differences in the number of DGRs or their taxonomic distribution (Supplemental Figure S7B-E). Together, these results demonstrate that mode of delivery has a significant impact on the number and types of microbes harboring DGRs, with vaginally born infants acquiring a larger number that more closely resemble the phylogenetic distribution observed in adults.

To determine if DGRs were active at any point within these samples, we aligned raw sequencing reads with identified VRs and searched for adenine-specific mutations. The percentage of active DGRs in infants varied from 56% to 80% over the first year of life but was similar to maternal and adult levels (Figure 6F). Next, we calculated the consensus VR amino acid sequence at each timepoint and compared VR haplotypes between mothers and infants. Of 388 transmitted DGRs, 72% showed evidence of VR haplotype switching (Figure 6G), raising the possibility that new variable proteins had been selected in the infant. A representative example of a diversified type V pilin homolog that was present in a mother at birth, and subsequently detected in her infant’s profile over the first year of life, is illustrated in Figure 6H. Comparing VR sequences shows that a majority of variable codons underwent continued alterations that changed the chemical class of diversified residues at four months, stabilizing by 1 year.

DGRs that were transmitted from mothers to infants born vaginally were most likely to be found in Bacteroidota (56%), rather than Bacillota (15%) (Figure 6E), and they were more likely to diversify pilus proteins than the general population of DGRs (15% *vs.* 10%, Chi-Square p<0.0001). Importantly, we were able to identify DGRs in our dataset that were nearly identical to each of the *Bacteroides* elements depicted in Figure 2B-D. Although the *Bfr, Bth,* and *Bun* DGRs we examined were quiescent under laboratory conditions (Figure 4B), homologous elements were highly active in mothers and infants, showing clear evidence of adenine-templated VR mutagenesis (Supplemental Figure S7F-G, Supplemental Table S15). For example, Figure 6I shows TR and VR sequences from a *Bun* DGR homolog that diversifies a Type V pilus tip adhesin, and the predominant VR haplotypes generated by a DGR that was transferred from mother to infant. As expected, elements homologous to the *Bov* and *Bfi* DGRs that were highly active *in vitro* and in germ-free mice were similarly active in humans.

These observations begin to characterize the abundance, distribution, and activities of DGRs in human infant and adult populations and their ability to diversify a wide array of potential ligand binding proteins, including pilus-associated adhesins. We also provide evidence that DGRs are transferred during the intergenerational handoff, that mode of delivery profoundly influences the relative abundance of DGR-containing microbes in the newborn gut, and that DGR transfer is associated with the appearance of new VR haplotypes that predominate in the infant’s gastrointestinal tract following maternal transmission, as would be expected for genotypic alterations that are subject to positive selection.

## Discussion

Numerous studies have highlighted the impact of microbial genetic variation on human health and disease^88–90^. This variation is driven by rapid adaptations that can quickly spread throughout microbial communities^91–93^. DGRs play a unique role in variation and adaptation by creating hypermutable hotspots that produce unrivaled levels of protein diversity, through a mechanism that targets ligand-binding residues and is shared across vast phylogenetic distances. By characterizing the variable proteins, mutational dynamics, and modes of transmission of *Bacteroides* DGRs, we can build a foundation for understanding how genome plasticity shapes host-associated microbial communities.

Of over 1,100 unique, DGR encoded variable proteins identified in human gut-associated *Bacteroides*, nearly 25% are predicted to diversify Type V pilins^48,63^. The genetic systems responsible for these structures display a remarkable degree of modularity and apparent redundancy, with the number, location, and organization of genes encoding pilin homologs differing between human-associated species and strains^48,63^. Less than half of the 16 multigene operons that encode pilin subunits in *Bfr* show evidence of protein expression *in vitro* (Supplemental Figure S2A), and many of these gene clusters are flanked by integrases, transposases, prophage loci, transfer genes and other signs of mobility^48^. Thus, differential expression and horizontal transfer may partly explain the modular complexity observed in *Bacteroides* genomes. For DGRs that target adhesive pilins in *Bfr*, *Bth*, and related taxa with homologous ICEs, we propose that two factors promote their dissemination and positive selection. The first involves properties of the conjugative elements, such as inducibility in mucus and the presence of stalk and anchor homologs that could help display diversified pilin tips. The second is the availability of new hosts with genetic backgrounds that are diverse and adapted to acquire horizontally transferred type V pilus genes. Adhesive pili are often observed to be essential determinants of microbe-host interactions involved in colonization, and microbe-microbe interactions that structure bacterial communities and facilitate biofilm formation^59,60^. An understanding of the selective advantages conferred by type V pili will benefit from the construction of isogenic mutants that are completely devoid of surface pili or lack specific components, the availability of animal models that recapitulate adhesive interactions in the human GI tract, and the identification of ligands recognized by static as well as DGR-diversified tip adhesins.

While the abundance of DGRs in nature attests to their selective advantages^12,19,55^, it is unknown how host cells balance the benefits of accelerated evolution with the increased potential for loss of fitness. It is reasonable to expect that mutagenic retrohoming will often be subject to regulation, with bursts of mutagenesis strategically deployed during times of stress, population expansion, nutritional changes or other factors, and interspersed with periods of quiescence that allow selection and fixation of adaptive traits^19^. Considering this, it was not unexpected to observe minimal activity with the *Bfr*, *Bth* and *Bun* DGRs *in vitro* or in gnotobiotic mice. In contrast, the constitutively high levels of activity measured with the *Bov* and *Bfi* elements was surprising, and provided an opportunity to examine DGR mutagenesis at a level of resolution that had not been previously attained. We observed an overwhelming (>1,000-fold) preference for introducing nonsynonymous substitutions in VR, a misincorporation bias that favors changes in the chemical properties of side chains available for ligand binding, and a role for host-encoded MutS that supports an all-or-nothing model for repair of heteroduplex intermediates during mutagenic retrohoming. In GF mice, we measured a decrease in the Shannon entropy of *Bov* VR sequences that was dependent on the presence of competing microbes and resulted in the appearance of new predominant VR haplotypes, as expected for conditions that favor positive selective sweeps.

Although signals and regulatory mechanisms that control mutagenic retrohoming await discovery, close homologs of all of the DGRs we studied were identified in human metagenomes and observed to be active in the human gut. This highlights the importance of examining these elements in their natural context. For DGRs predicted to diversify bacterial factors, type V pilus subunits were the most common variable proteins encoded by Bacteroidota, and they were the most likely DGRs to show evidence of maternal-infant transfer. Vaginally born infants had a greater number of DGRs compared to infants born by C-section, and they were mainly encoded by *Bacteroides* species as opposed to Bacillota. This difference resolved quickly and mirrors well-characterized time-dependent effects of birth-mode on the overall composition of the developing infant microbiome^94^. Most interesting, however, is that for the majority of variable proteins identified as being transferred, the predominant VR haplotypes observed in mothers switched to new predominant haplotypes in their infants, consistent with the hypothesis that DGR-driven adaptations are occurring during the first year of life.

Hypervariable systems and assessments of purifying selection may provide a means for identifying genes and networks in microbial communities that confer contextually significant fitness advantages. Accordingly, our working hypothesis is that DGRs that diversify bacterial proteins function, at least in part, to optimize colonization factors that promote engraftment and maintenance of host bacteria within the GI microbiota. If true, understanding DGRs and the variable proteins they diversify will not only have applications for understanding microbe-host interactions, but may also provide a means to engineer therapeutic microbial consortia capable of rapidly evolving colonization factors to promote efficient engraftment in new hosts. This could pave the way for future applications that harness the adaptive properties of DGRs to support health and reverse microbiome-associated diseases.

## Limitations

There are two material limitations of our study that reflect broader challenges in efforts to understand diversity in human-associated microbial communities. The first involves the need to identify phenotypes from genotypic information, a transformative resource available on a massive scale that is rarely sufficient to provide causal links. DGRs were discovered based on a phenotype, tropism switching by *Bordetella* phage, and the genetic basis was characterized with relative ease^17,18^. In contrast, identifying the function of an uncharacterized gene, or the advantage of diversifying it, may require recapitulating natural environments that provide appropriate selective pressures, including signals for expression, receptors for ligand interactions, and many other context-dependent parameters. The ‘genotype to phenotype to mechanism’ pathway can pose major challenges, but they are not necessarily insurmountable. In the case of DGRs and similar systems, for example, hypervariability can provide sensitive, real-time estimates of positive selection and the environmental conditions under which it occurs – information that can be used to target hypothesis-driven discovery.

A second limitation involves the nature of metagenomic data available from existing large-scale efforts to characterize human microbiomes, which is primarily derived from short read approaches that make assembly difficult and rarely reach sufficient depth to fully capture VR sequence variability and entropy. Future efforts that incorporate long-read sequencing of sufficient depth on well curated longitudinally obtained mother-infant samples, along with Amplicon-Seq to deeply characterize diversified VRs, will be required to understand DGR dynamics following birth and to identify variable proteins subject to positive selection in developing infants.

## Acknowledgements

We thank Elaine Hsiao and her laboratory members Kristie Yu and Jorge Paramo at UCLA, as well as Sarkis Mazmanian at the California Institute of Technology for valuable advice and assistance with gnotobiotic mouse husbandry. We also thank Eric Martens at the University of Michigan for providing several of the *Bacteroides* strains used in this study. We are indebted to Suzanne Devkota at Cedars Sinai Medical Center, Partho Ghosh at the University of California, San Diego, and Blair Paul at the Marine Biological Laboratory, Woods Hole, for their insightful critiques of the manuscript. This research was supported by the NIH (1K08DK138316/5K12HD111040; 5K12HD000850 to B.R.M.) and the JDS Family Foundation and Kavli Endowment (to J.F.M.). We acknowledge the use of resources at the UCLA Proteomics Laboratory for mass spectroscopy and the UCLA Neuroscience Genomics Core for RNA-sequencing. The funders had no role in the design of the study, in the collection, analyses, or interpretation of data, in the writing of the manuscript, or in the decision to publish the results.

## Author contributions

Conceptualization, B.R.M., Y.W., U.A., and J.F.M, with input from all authors; Methodology, B.R.M, Y.W., U.A., and J.F.M; Investigation, B.R.M., Y.W., C.W., J.R., K.S., C.B., S.M., and U.A.; Software, B.R.M. and B.M.S.; Writing – Original Draft, B.R.M. and J.F.M.; Writing – Review and Editing, B.R.M., U.A., and J.F.M.; Funding acquisition, B.R.M. and J.F.M;

## Declaration of interests

J.F.M. is a cofounder and chair of the scientific advisory board (SAB) of Pylum Biosciences, Inc., a member of the SAB of Notitia Biotechnologies, and an advisory board member of Seed Health, INC.

## Supplemental information

Figures S1–S7.

Excel file containing Tables S1-S15.

**Supplemental Figure S1.**
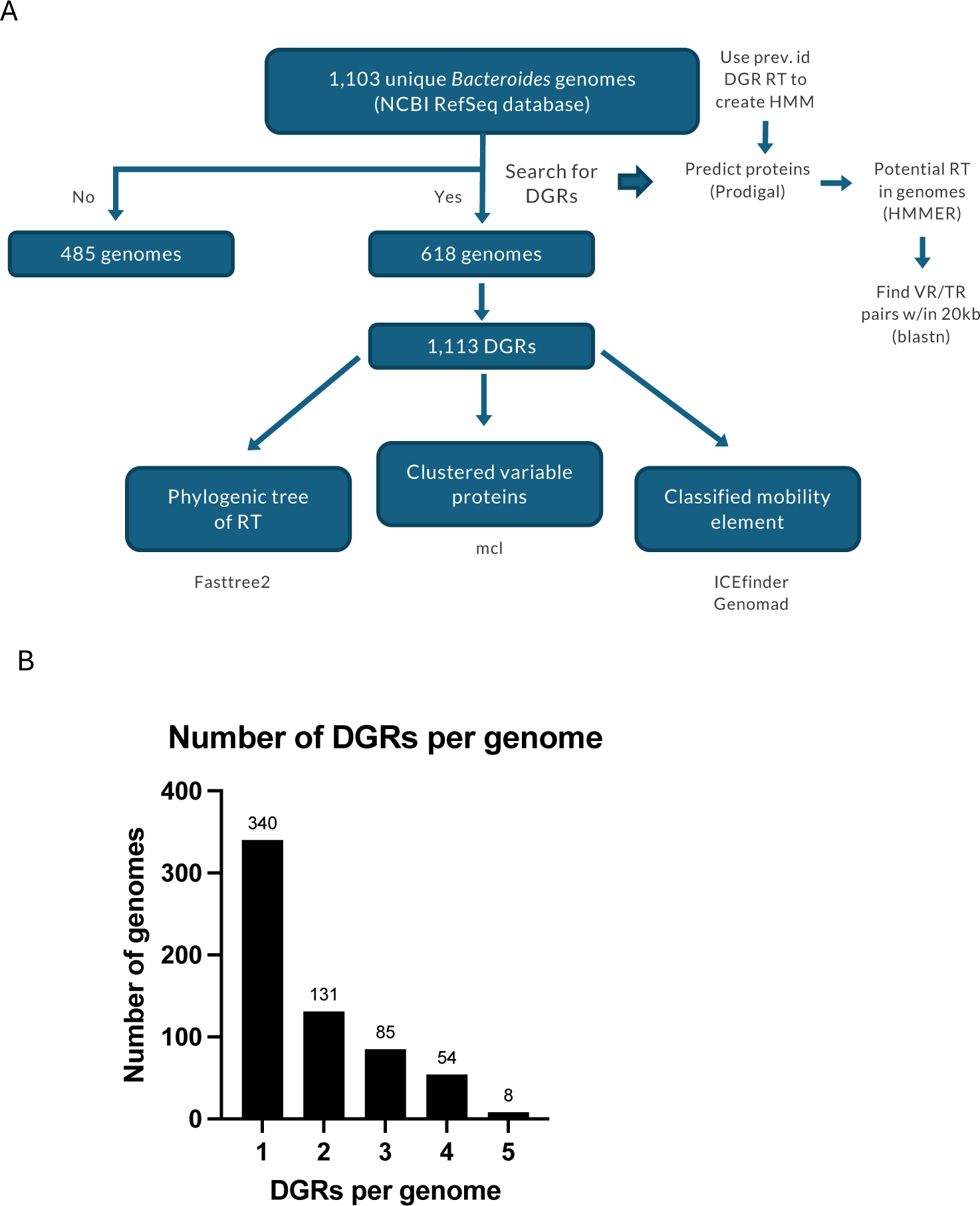
Overview of *Bacteroides* DGRs, related to Figure 1: (A) Schematic of the DGR identification methodology from *Bacteroides* genomes. Proteomes were first predicted from assemblies using Prodigal^1^ and were used as input for profile-based searches for the DGR RT proteins using HMMER^2^. Imperfect repeats representing VR and TR were searched within a 20 kb window upstream and downstream of potential *rt* genes using blast^3^. Imperfect repeats were further filtered across two criteria. First, pairs had to differ from each other at positions that correspond to adenines in one of the repeats. Second, the predicted VR repeat had to be located within a gene encoding region previously predicted by Prodigal. Using the identified DGRs, a phylogenetic tree was built using FastTree2^4^, the variable proteins were clustered using blastp^3^ and mcl^5^. GeNomad^6^ and ICEBerg v2^7^ were used to determine if the DGR locus fell within an ICE or prophage genome. (B) Distribution of the number of DGRs identified within the same genome across all *Bacteroides* strains in our dataset.

**Supplemental Figure S2.**
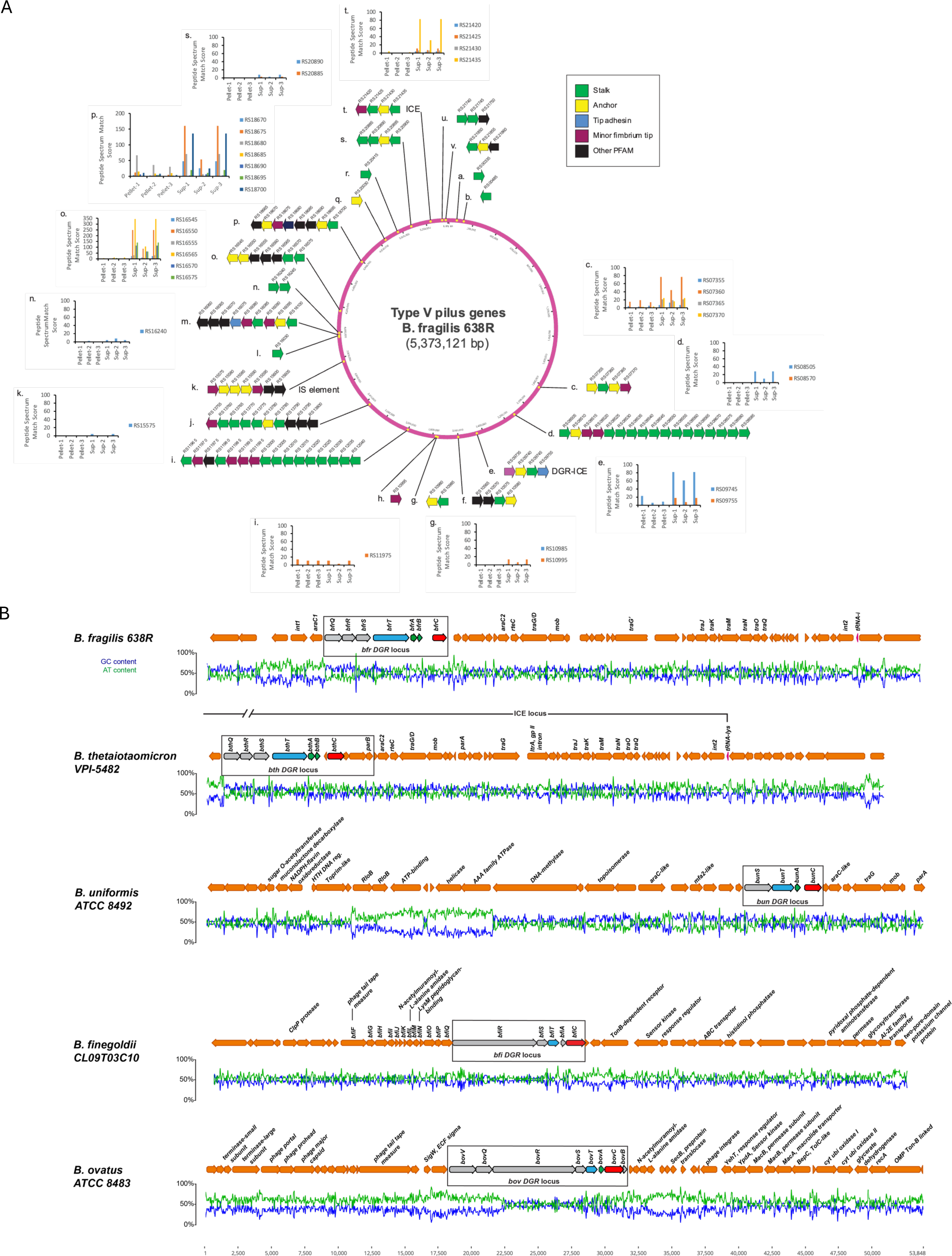
*Bacteroides* pilin loci, related to Figure 2: (A) Overview of the 22 loci within the *Bfr* genome that encode pilus proteins. Each gene was functionally categorized according to its homology to known pilus proteins using HHpred^8,9^. Bar graphs for the indicated loci display the Peptide Spectrum Match Score across individual proteins identified by mass spectroscopy of *Bfr* cells grown *in vitro*. (B) Genomic loci and GC content adjacent to DGRs in the *Bacteroides* strains indicated. Within each DGR locus, the target gene is colored blue, *rt* is red, and the accessory genes are green. Predicted DGRs with accessory genes (colored gray) are boxed. Genes with predicted annotations are indicated with the name of the gene. Genes without predicted names have unknown function or are hypothetical.

**Supplemental Figure S3.**
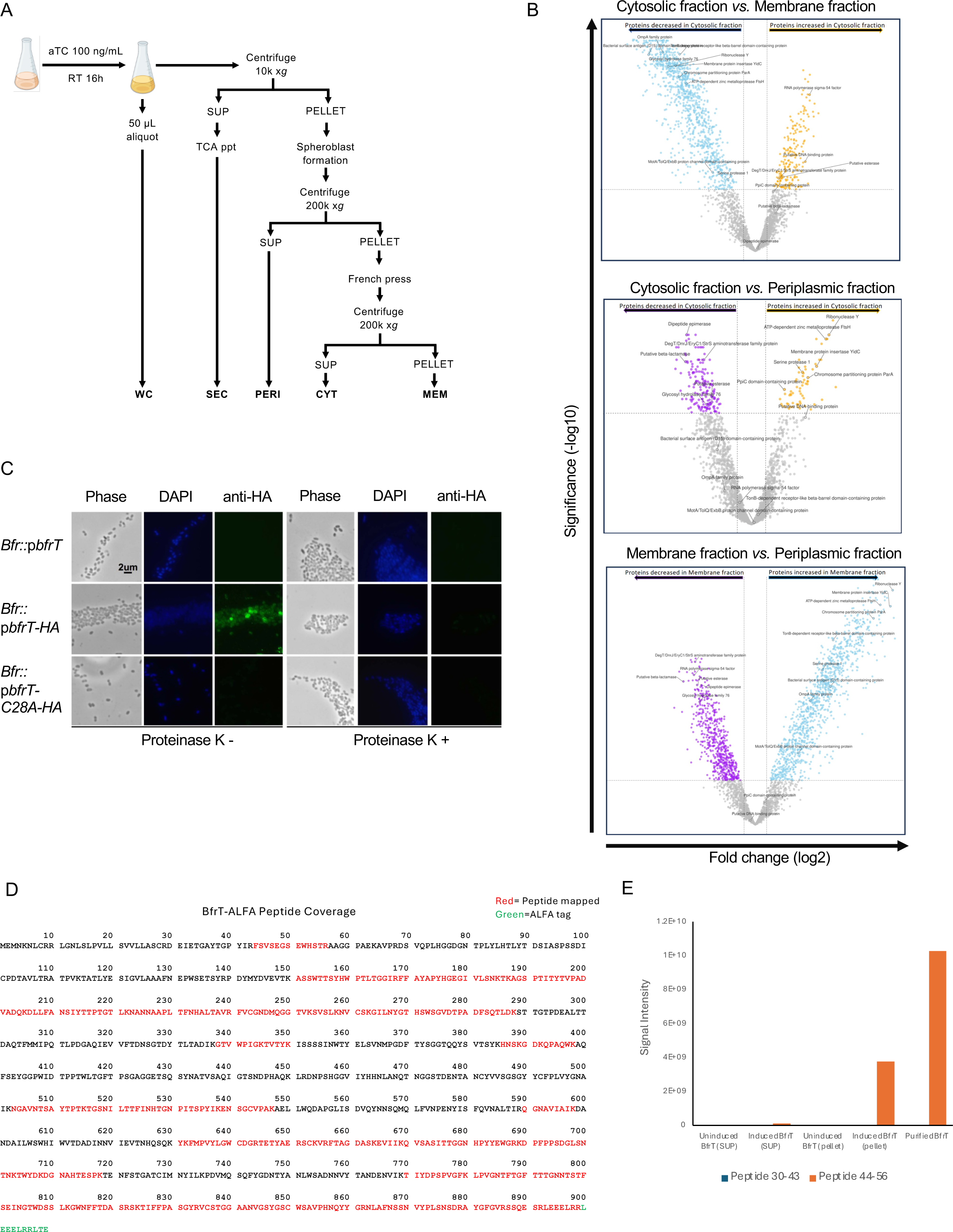
*Bacteroides* diversify pilus proteins, related to Figure 2. (A) Schematic of cellular fractionation methodology. Cells were either induced with 100 ng/mL of aTC or not and incubated at room temperature for 16 hours. A 50 μL aliquot was set aside for the whole cell fraction. The remaining culture was centrifuged at 10,000 x g. The supernatant was TCA precipitated and stored as the secreted fraction. The pellet fraction was subjected to spheroblasting and centrifuged at 200,000 x g. The resulting supernatant was stored as the periplasmic fraction while the pelleted fraction was passed through a French press to separate cytoplasmic and crude membrane fractions. (B) Volcano plot of relative protein enrichment between cellular fractions in *Bov* cells grown *in vitro*. The abundance of individual proteins from the periplasmic, cytosolic, and membrane fractions was quantified by mass spectrometry and the relative abundance change of each protein was calculated for each fraction comparison (i.e., cytosol fraction compared to membrane fraction; cytosol fraction compared to periplasmic fraction; membrane fraction compared to periplasmic fraction). The VolcaNoseR^10^ web application was used to plot the Log_2_ fold change of protein abundance versus the -Log_10_ of the Significance value of the change in abundance for each protein between each of the fractions. Specific proteins with previously reported sub-cellular localization data were tracked to evaluate the effectiveness of the fractionation protocol (see Supplemental Table S8). (C) Immunofluorescence of *Bfr* cells expressing epitope tagged BfrT. Cells overexpressing WT BfrT (untagged), HA-tagged BfrT, or the HA-tagged BfrT-C28A mutant were treated with or without proteinase K. (D) Summary of the peptides identified by mass spectroscopy that aligned to BfrT from the supernatant of *Bfr* cells grown *in vitro*. (E) Signal intensity of the peptide fragments recovered from Part D for peptides 30-43 and 44-56, which represent arginine cleavage sites during pilus assembly.

**Supplemental Figure S4.**
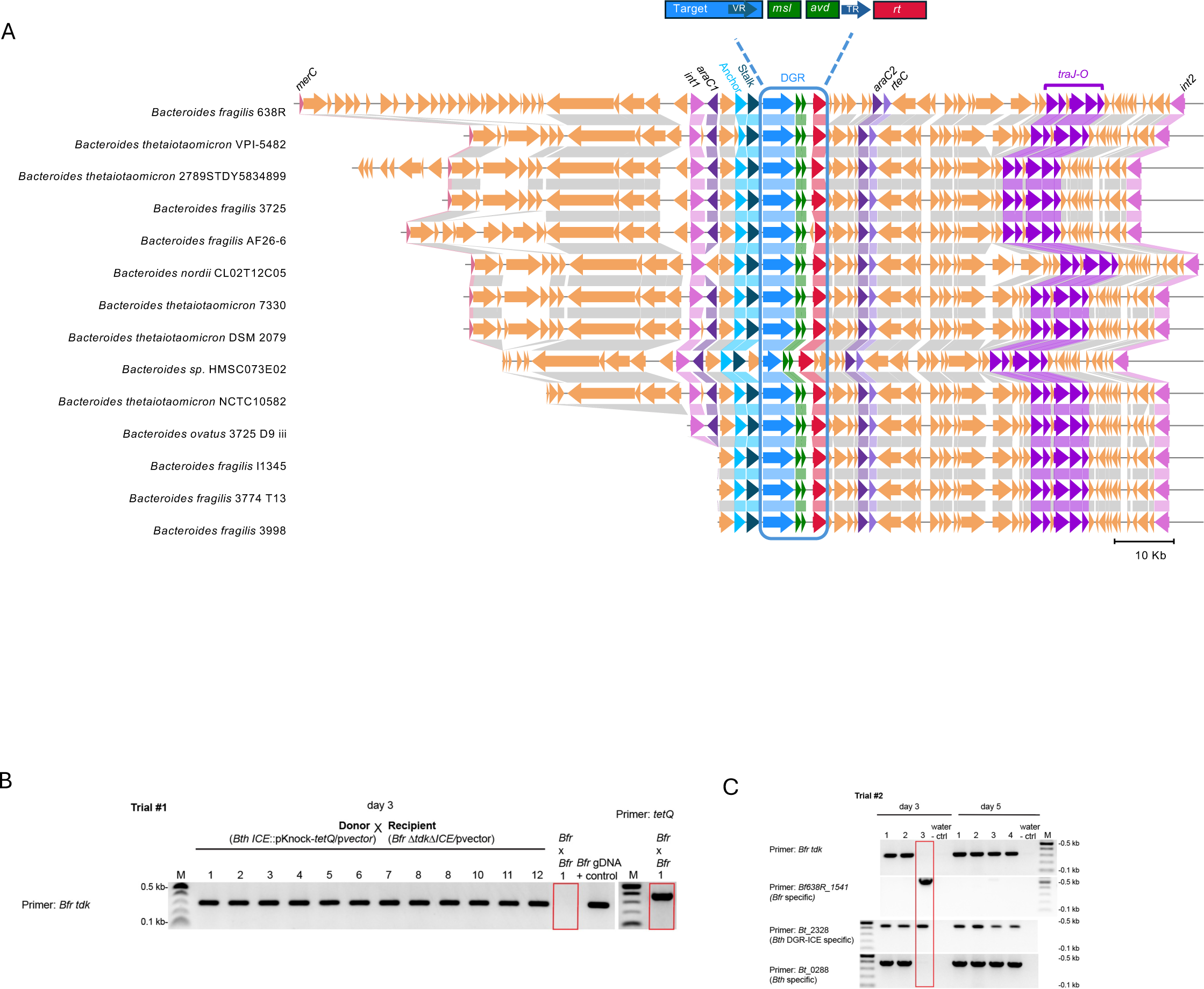
Overview of *Bacteroides* ICEs, related to Figure 3. (A) Synteny of 16 DGR-encoding ICEs identified in different *Bacteroides* strains. The name of the *Bfr* gene is labeled on top for genes with predicted functions. Genes are connected if their encoded proteins have at least 90% identity. The DGR locus within the ICEs is boxed. (B) PCR based screen to identify ICE transconjugants from matings between *Bfr* and *Bth* cells. True transconjugants, as shown by the red box, are negative for the *tdk* gene while positive for the ICE (*tetQ*). (C) PCR based screen as in Part B but using *Bth* cells as donor cells.

**Supplemental Figure S5.**
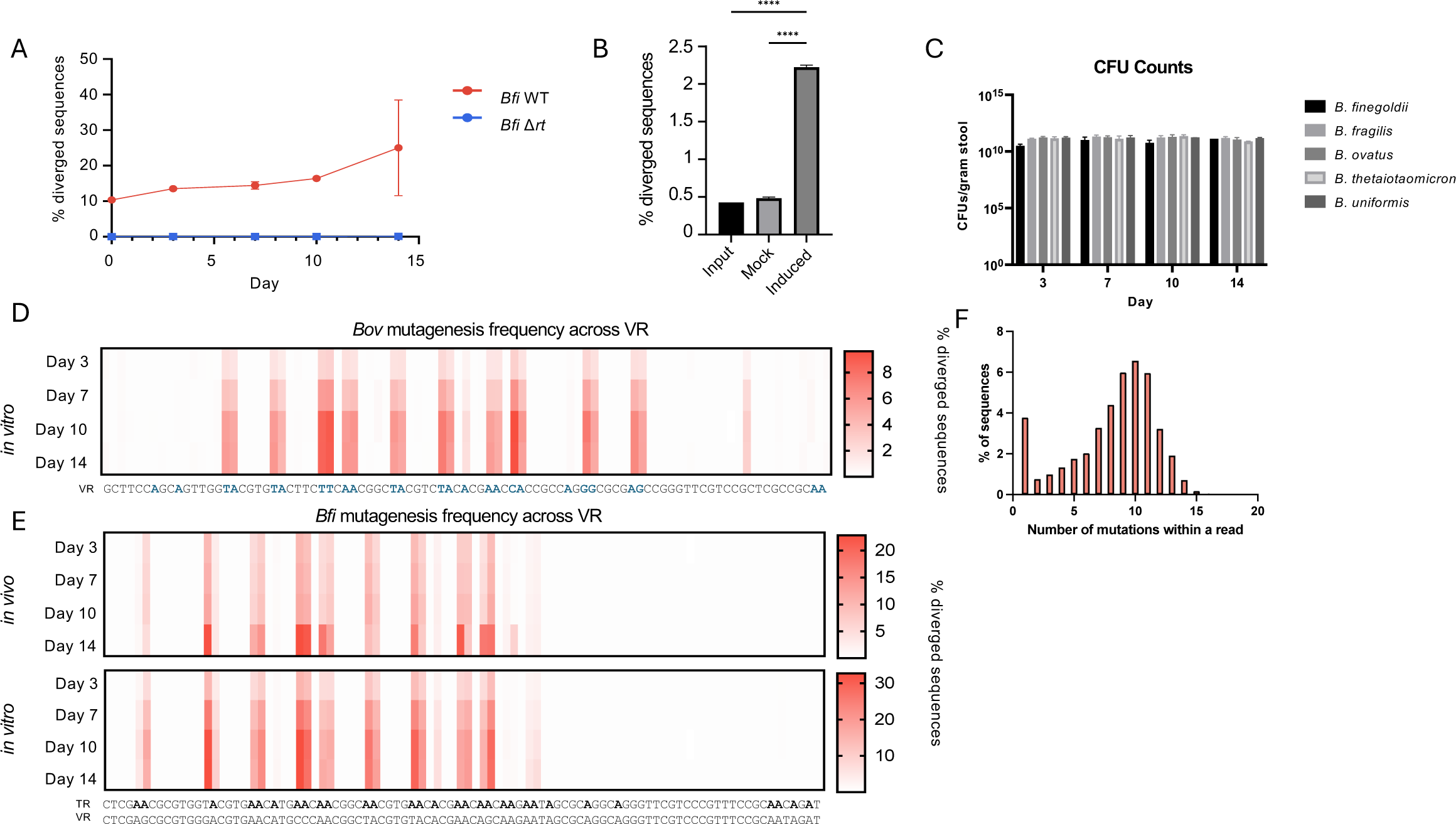
DGRs are active in *Bacteroides*, related to Figure 4. (A) Percent of VRs that diverged from their cognate parental VR in *Bfi* WT and *Bfi* Δ*rt* present in fecal samples from monocolonized SW mice (n=4 each). Error bars, standard deviation of mean. (B) Number of VR reads that diverged from their cognate parental VR sequence in *BovΔrt::pNBU2-tetR* P1T_DP_^GH023^-*rt*, a mutant in which the chromosomal copy of the DGR *rt* has been knocked out and WT *rt* is expressed ectopically under the control of an inducible promoter. Mock induction or induction of *rt* transcription with aTC are shown. Error bars, standard deviation from mean. ****p<0.0001, ANOVA. (C) Number of CFUs obtained per gram of stool from monocolonized SW mice over a two-week period (n=4); error bars, standard deviation from mean. All comparisons across strains are non-significant (n.s.). (D) Mutation frequencies of individual nucleotide positions within the *Bov* target gene over a two-week period from cells grown *in vitro*. (E) Mutation frequences of individual nucleotide positions within the *Bfi* target gene over a two-week period present in fecal samples from monocolonized SW mice (top) or from cells grown *in vitro* (bottom). (F) Distribution of the number of nucleotide substitutions per VR read in *Bfi* cells grown *in vitro* at Day 14.

**Supplemental Figure S6.**
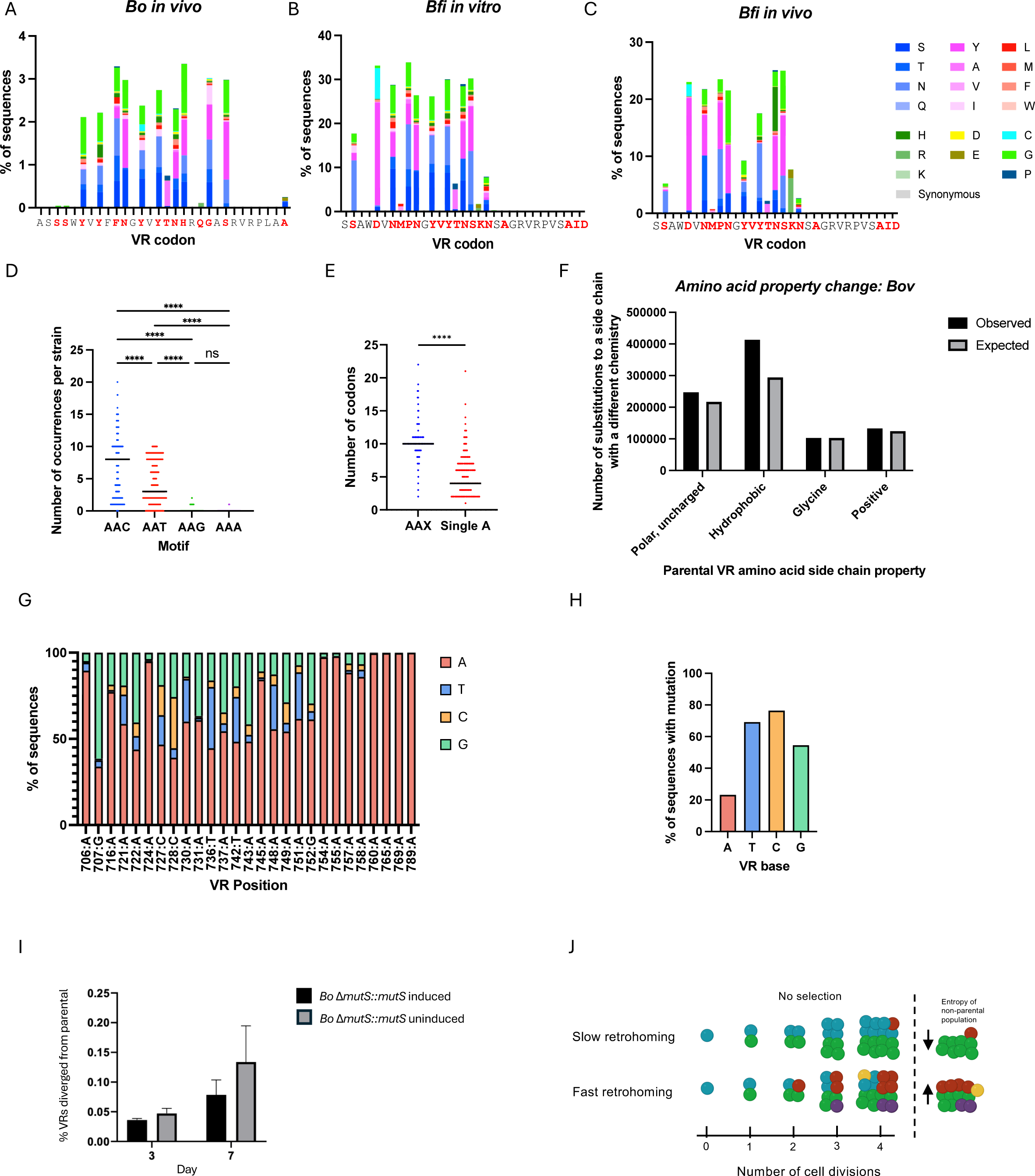
*Bacteroides* DGRs preferentially create non-synonymous mutations, related to Figure 5. (A) Substitution frequency at *Bov* VR codons present in fecal pellets of monocolonized SW mice at 14 days post-gavage. (B) Substitution frequency at *Bfi* VR codons from cells grown *in vitro* after 14 days. (C) Substitution frequency at *Bfi* VR codons present in fecal samples of monocolonized SW mice at 14 days post-gavage. (D) Number of AAX motifs present in *Bacteroides* TRs. ****p<0.0001, ANOVA, Holm-Sidak multiple comparison correction. N.s, not significant by ANOVA test. (E) Number of AAX motifs vs motifs containing a single adenine in *Bacteroides* TRs. **** p<0.0001, Mann-Whitney test. (F) Number of occurrences that a diversified codon would encode for an amino acid with a different chemical property of its side chain from *Bov* cells grown *in vitro*. The expected number was calculated under the assumption that every AAX motif has an equal chance of occurring through mutagenic retrohoming. (G) Nucleotide frequency at individual variable VR positions within diversified *Bfi* VRs. The parental VR position and nucleotide are listed under each bar. (H) Cumulative frequency of each of the four nucleotides at variable VR sites within diversified *Bfi* VRs. (I) Percent of VRs that diverged from their cognate parental VR in *BovΔmutS::pNBU2-tetR* P1T_DP_^GH023^-*mutS,* a mutant in which the chromosomal copy of *mutS* was knocked out and WT *mutS* was expressed ectoptically under an aTC-inducible promoter, following aTC induction or mock induction (n=3). Error bars, standard deviation from mean. (J) Representation of the effect on VR diversity of slow mutagenic retrohoming *vs*. fast mutagenic retrohoming with respect to cell division. Blue spheres represent cells with the parental VR sequence, all other colored spheres represent cells with diversified VR sequences.

**Supplemental Figure S7.**
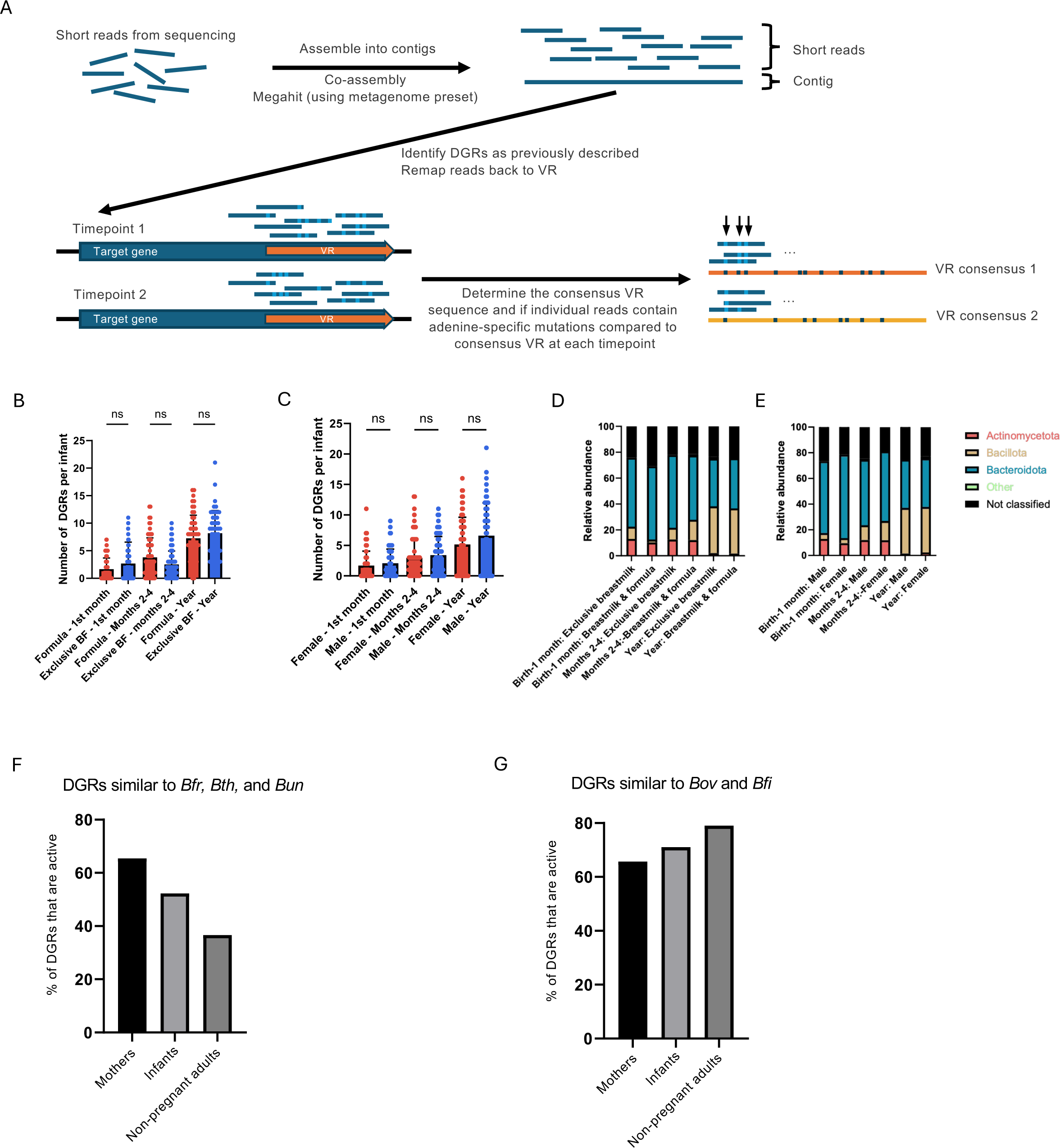
DGRs in mother-infant datasets, related to Figure 6. (A) Schematic of the methodology for DGR identification and activity detection from metagenomic sequencing reads. (B) Number of DGRs identified per infant, grouped by feeding preference and age. n.s. not significant by ANOVA tests. (C) Number of DGRs identified per infant, grouped by gender and age. n.s. not significant. (D) Taxonomic distribution of DGR-containing microbes at different ages, grouped by feeding preference. (E) Taxonomic distribution of DGR-containing microbes at different ages, grouped by gender. (F) Percent of active DGRs with target genes similar to the target genes of *Bfr, Bth,* and *Bun* in mothers, infants, and healthy adults. (G) Percent of active DGRs with target genes similar to the target genes of *Bov* and *Bfi* in mothers, infants, and healthy adults.

## Methods

### Bacterial strains, plasmids, primers, and growth conditions

Bacterial strains are described in Supplementary Table 1, plasmids are described in Supplementary Table 2, and primers are described in Supplementary Table 3. *E. coli* strains were grown aerobically at 37°C in Luria-Bertani medium (LB, BD Difco). *Bacteroides* strains were grown in Brain Heart Infusion (BHI, BD Difco) medium at 37°C, supplemented with vitamin K (5 µg/mL) and hemin (5 µg/mL), and incubated in an anaerobic chamber (Coy) under 5% H_2_, 10% CO_2_, 85% N_2_. Antibiotics were added to the following final concentrations: Carbenicillin (Cb), 100 μg/mL; Gentamicin (Gm), 200 μg/mL; Erythromycin (Erm), 5 or 25 μg/mL, Tetracycline (Tet), 2 μg/mL, 5-fluoro-2’-deoxyuridine (FudR, 200 μg/mL), unless otherwise noted. Anhydrous tetracycline (aTC) was used at a concentration of 100 ng/mL. Backbone plasmids used in this study were pLGB13, pLGB36, and pNBU2_erm-TetR-P1T_DP-GH023^102–104^.

### Mice

Germ-free Swiss Webster mice were purchased from Taconic Farms and bred in flexible film isolators. For gnotobiotic experiments, sterile litters of 8-10 week old male and female mice were transferred into autoclaved microisolator cages where they were fed autoclaved chow diets *ad libitum* and given autoclaved water supplemented with gentamicin 100 μg/mL. Altered Schaedler Flora (ASF) live animal donor C57BL/6 mice were purchased from Taconic (ASF-DONOR-M/F) and fed an autoclaved chow diet *ad libitum* and given autoclaved water. For monocolonization experiments, animals were orally gavaged with a 200 μL inoculum of an overnight culture of a given *Bacteroides* strain containing ∼10^8^ cells per gavage. For ASF experiments, approximately 1 g of fresh stool from an ASF donor mouse was vortexed into 1 mL of an overnight culture of *Bacteroides* immediately prior to oral gavage. For conjugation experiments, 100 μL of each donor and recipient strains were combined in a 1:1 ratio and orally gavaged to mice. Stool samples were collected on Days 3, 7, 10, and 14. All procedures were performed in accordance with an approved protocol following IUCAC guidelines at the University of California, Los Angeles and the IUCAC guidelines at the California Institute of Technology.

### Plasmid generation

Q5 high fidelity DNA polymerase (New England Biolabs [NEB]) or Phusion high fidelity DNA polymerase (NEB) were used for PCR cloning steps. To construct plasmids for deletion mutagenesis, a PCR fragment was generated that included 1kb upstream and the first 12 codons of the targeted gene and a second fragment was generated to include the last 12 codons and 1kb downstream of the targeted gene. The backbone plasmids pLGB36 (*Bfr* only) or pLGB13 (all others) were cut at the BamHI (NEB) restriction site. PCR fragments were ligated together and into the backbone plasmid using NEBuilder HiFi assembly master mix (NEB). For overexpression mutants, the targeted gene was cloned into pNBU2_erm-TetR-P1T_DP-GH023 plasmid cut at NcoI and SalI restriction sites using the NEBuilder HiFi assembly master mix. Plasmids were sequenced following cloning to ensure insertions were without mutations.

### Conjugation and allelic exchange

Plasmids were transformed into *E. coli* S17-λpir and conjugated into *Bacteroides* strains as previously described^102^. Briefly, *E. coli* S17-λpir donor cultures were grown in 25 mL of LB+Cb to an OD_600_ between 0.3-0.5. Recipient *Bacteroides* strains were grown in BHIS to an OD_600_ of 0.05-0.1. Both strains were mixed together in a 50 mL Falcon centrifuge tube and spun at 4000x*g* for 10 mins. Cell pellets were then resuspended in 100 µL of BHIS, plated on the center of a prewarmed BHIS plate, and incubated aerobically at 37°C for 14-16 hours. The mating spot was resuspended in BHIS and was diluted serially from 1:10 to 1:10,000. From each dilution, 100 µL was streaked onto BHIS+Gm+Erm selection plates and incubated at 37°C anaerobically. Colonies were picked after two days, restreaked on BHIS+Gm+Erm plates, and grown for two additional days. For overexpression mutants, stocks were made from an overnight culture of BHIS and integration was confirmed by PCR of attB sites using the appropriate primers in Supplemental Table 3. For allelic exchange protocols, isolates were grown in BHIS overnight, diluted to 1:1,000 or 1:10,000 in the morning, and plated on BHIS+aTC100 to induce the counterselection toxin. After two days, colonies were picked, and colony PCR screening was performed to determine which colonies contained the desired mutation and which colonies reverted back to WT. Stocks were made of potential mutants. The mutation site including 1kb upstream and downstream of the mutation, as well as the entire DGR locus was amplified from chromosomal DNA and sequenced to ensure no undesired mutations were introduced.

### Sample preparation and immunoblotting

For SDS-polyacrylamide gel electrophoresis (SDS-PAGE) sample preparation, *Bacteroides* strains were cultured in BHIS media and harvested as described previously^105^. Proteins with ALFA-tags were detected using a monoclonal mouse antibody at a dilution of 1:5000 (anti-ALFA, NanoTag Biotechnologies). Immunodetection was carried out by chemifluorescence using horseradish peroxidase-labelled goat anti-mouse IgG and the ECL plus^®^ detection substrate (GE Healthcare). Chemifluorescent signals were visualized using a Typhoon scanner (GE Healthcare).

### Cellular fractionation

Cellular fractionation was carried out as described^106^ and is briefly summarized here. Overnight cultures of WT or mutant *Bacteroides* strains encoding inducible ALFA-tagged proteins were diluted 1:100 in a total volume of 1 L BHIS supplemented with 25 µg/ml Erm and incubated at 37°C until reaching OD 0.05-0.1. The target gene was then induced by addition of 100 ng/mL of aTC to the media and the culture was allowed to incubate for an additional 16 hr at RT. The induced 1 L culture was then pelleted by centrifugation at 10,000xg and subsequently resuspended in 10 mL of spheroblast buffer (0.2 M Tris-HCl pH 8, 1 M sucrose, 1 mM EDTA, 1 mg/mL lysozyme) and incubated for 5 min at RT. A volume of 40 mL of ice-cold dH2O was then added to the suspension before placing on ice for 5 min to allow spheroblast formation. The suspension was then centrifuged at 200,000xg for 45 min at 4°C. The resulting supernatant was collected as the periplasmic fraction and the pellet was resuspended in French press buffer (7.5 mL ice-cold 10 mM Tris-HCl pH 7.5, 5 mM EDTA, 0.2 mM DTT, 50 μL 1mg/mL DNaseI). Cells were then ruptured in a French Press with two passes at 10^8^ Pa. Unbroken cells in the lysate were removed by centrifugation at 10,000xg for 10 min at 4°C. The lysate was then centrifuged at 280,000xg for 4hr at 4°C. The resulting supernatant was collected as the cytoplasmic fraction while the pellet contained crude membranes. Membrane fractions were diluted 1:1 with dH2O, centrifuged at ≥85 000xg for 20 min at 4°C, washed 3x in 500 μL dH2O and then stored at −20°C.

### Proteinase K

Overnight cultures of *Bacteroides* strains carrying an inducible, ALFA-tagged variable protein (*Bfr* WT-ALFA, *Bfr* C28A-ALFA, or *Bov* WT-ALFA) were diluted 1:100 in a total volume of 10 mL BHIS supplemented with 25 µg/ml Erm and incubated at 37°C until reaching OD 0.1. Expression of the ALFA-tagged variable protein was then induced by addition of 10 ng/mL aTC. Induced cultures were incubated for 8 hrs and then collected by centrifugation. Cells were resuspended at 2.5 OD/mL in a total volume of 5 mL PBS and 1 mL aliquots of the bacterial slurry were dispensed to 1.5 ml tubes and incubated for 1 hour at 37°C with one of the following Proteinase K quantities : 0 (control); 25 ng; 50 ng; 100 ng; 200 ng. Next, 10 ul PMSF was added to each tube and cells were collected by centrifugation at 10,000 xg and washed 2X with 1 mL PBS. Cells were resuspended in SDS sample dye with beta-mercaptoethanol, boiled for 10 minutes and analyzed by Western blot.

### ALFA pulldown

ALFA-tagged *Bacteroides* variable proteins and their interacting protein partners were purified from an appropriate cellular fraction (periplasm for *Bov*, supernatant for *Bfr*) using the Anti-ALFA single domain nanobody resin (ALFA SelectorST) according to manufacturer instructions. Briefly, 3 mL of the cellular fraction was diluted 1:1 with 2x binding buffer (50 mM Tris pH 7.5, 100 mM NaCl, 2 mM EDTA, 1 % NP-40, 10% glycerol) supplemented with 10 uL/ml HALT protease (ThermoFisher). To this suspension, 200 μL of ALFA SelectorST was added and the mixture was incubated at 4°C with end-over-end rotation for 16 hr. The resin was collected by centrifugation and washed 2x with PBS containing 0.5% NP-40. The resin was then pelleted and resuspended in 200 μL of 1x Laemmli buffer (0.0625 M Tris pH 6.8, 2% sodium dodecyl sulfate, 10% glycerol, bromphenol blue) with 1% β-mercaptoethanol and then boiled for 5 min.

### Mass spectroscopy

Immunoprecipitation eluates (ALFA pulldowns) or cellular fractions (fractionation controls) in 1x Laemmli buffer were diluted in equal volume of 100mM Tris-Cl pH 8.5 and reduced and alkylated by the sequential addition of 5 mM tris(2-carboxyethyl) phosphine and 10 mM iodoacetamide. This was followed by treatment with single-pot, solid-phase-enhanced sample preparation (SP3) protocol for protein clean-up^107^. Following SP3, eluates were proteolytically digested with Lys-C and trypsin at 37°C overnight. The digested peptides were subjected to offline SP3-based peptide clean-up and subsequently analyzed by LC-MS/MS. Briefly, peptides were separated by reversed-phase chromatography using 75 μm inner diameter fritted fused silica capillary column packed in-house to a length of 25 cm with bulk 1.9 mM ReproSil-Pur beads with 120 Å pores. The increasing gradient of acetonitrile was delivered by a Dionex Ultimate 3000 (Thermo Scientific) at a flow rate of 200 nL/min. MS/MS spectra were collected using data-dependent acquisition on an Orbitrap Fusion Lumos Tribrid mass spectrometer (Thermo Fisher Scientific) with an MS1 resolution (r) of 120,000 followed by sequential MS2 scans at a resolution (r) of 15,000. The data generated by LC-MS/MS were analyzed using the MaxQuant bioinformatic pipeline^108^. The Andromeda integrated in MaxQuant was employed as the peptide search engine. Briefly, a maximum of two missed cleavages was allowed. The maximum false discovery rate for peptide and protein was specified as 0.01. Label-free quantification (LFQ) was enabled with LFQ minimum ratio count of 1. The parent and peptide ion search tolerances were set as 20 and 4.5 ppm respectively. The MaxQuant output files were subsequently processed for statistical analysis of differentially enriched proteins using Analytical R tools for mass spectrometry (artMS)^109^.

### Cellular Fractionation Controls

To assess the quality of protein enrichment from our cell fractionation protocol, we used mass spectrometry to identify all proteins in each fraction generated and then compared the change in the abundance of each identified protein between each fraction (i.e., cytosol fraction compared to membrane fraction; cytosol fraction compared to periplasmic fraction; membrane fraction compared to periplasmic fraction). For each protein, the change in its abundance between fractions was graphed using the VolcaNoseR^110^ web application. The log_2_ fold change of abundance was graphed over the x-axis and the -log_10_ significance value of the change in abundance was graphed over the y-axis. We then tracked specific proteins with previously reported sub-cellular localization data to evaluate the effectiveness of the fractionation protocol (Supplementary Table S8).

### Detection of DGR-ICE integration, excision, and episome formation by PCR

Genomic DNA was prepared from cultured *Bfr* or *Bth* cells carrying overexpression constructs (pEV, p*araC2*, p*rteC*, p*merR*) using a DNeasy Blood and Tissue Kit (Qiagen). A 50 ng aliquot of DNA was used as template in a standard 50 μL PCR with primer sets that detect integrated DGR-ICE junction fragments or episomes and chromosomal scars resulting from ICE excision (Figure 3A). To quantify ICE excision *in vivo*, germ-free Swiss Webster mice (n=3) were monocolonized by *Bfr* carrying the plasmid pFD340, which confers Erm resistance. Bacterial DNA was extracted from mouse feces and cecal content using the ZR fecal DNA miniprep kit (Zymo Research). DNA from *Bfr* cells in scraped colon mucus was prepared using DNeasy Blood and Tissue Kits (Qiagen) following pre-treatment with N-acetyl-L-cysteine. Specifically, a freshly made NALC solution (50ml 2.94% sodium citrate, 50ml 4% sodium hydroxide, 500mg N-acetyl-L-cysteine) was added to the mucus at 1:1 ratio (vol/vol), incubated at the room temperature for 1 hr with agitation until the sample attained desired fluidity. Quantitative PCR (qPCR) was performed on an iCycler iQ real-time PCR detection system (BioRad) with iQ SYBR Green Supermix (BioRad). Per each 30μl qPCR, 30 ng DNA from various samples was used as template. Ct values for the episome and chromosomal scar were normalized to Ct values of the housekeeping gene, rpoD (BF638R_RS13245). Relative quantification of excised ICE in different samples was calculated by the ΔΔCt approach, using cultured *Bfr* with pFD340 as baseline. DNA extracted from *in vitro* grown cultures of *Bfr* carrying p*araC2* was included as a positive control for high level excision.

### ICE Transfer assays

To create ICEs and strains with compatible antibiotic markers, the Erm resistance cassette in pKnock-erm was first swapped with a *tetQ* marker, resulting pKnock-tetQ. The DGR-ICE from *Bfr* or *Bth* was tagged with the Tet resistance marker by inserting pKnock-tetQ downstream of *rt*. Mating experiments were designed to measure inter- and intra-species ICE transfer. Briefly, three independent cultures of donor or recipient cells were grown to mid-log phase (OD_600_ 0.5), mixed at a 2:1 ratio and spotted on sterile nitrocellulose membranes (0.45 μM, PALL Life Science). After incubation on non-selective BHIS plates for 16-24 hrs, mating mixtures were washed off the filter into 2 mL of BHIS and plated onto selective BHIS medium with 2 μg/mL Tet and 200 μg/mL FudR. Putative transconjugants were purified and verified by PCR reactions with primer sets that are specific to the recipient or donor ICE. The transfer efficiency was calculated by dividing the number of genuine transconjugants by the number of recipients in each mating experiment. A similar transfer was also set up in gnotobiotic mice, except that both the tetQ-tagged donor and Δ*tdk*ΔICE recipient carry pFD340 to confer erythromycin resistance. Each mating pair was inoculated to 4 mice co-housed in one cage. Mouse feces were plated at days 1, 3, 5, 7 and 10 post-inoculation.

### Assays for mutagenic retrohoming

For *in vitro* assays, overnight cultures of *Bacteroides* strains were diluted to OD_600_ 0.01 in 3 mL of BHIS in triplicate. Every eight hours, the OD_600_ was measured (OD_600_ between 0.5-0.8), and cultures were rediluted to OD_600_ 0.01 in 3 mL of fresh BHIS. Samples were collected by pelleting 1 OD_600_ of cells on Days 3, 7, 10, and 14 for Amplicon-Seq. *In vivo* assays were conducted by collecting fecal pellets from monocolonized SW mice on Days 3, 7, 10, and 14 post-gavage. Genomic DNA was extracted from pellets for Amplicon-Seq.

### Amplicon-Seq of VR regions

Total DNA was extracted from bacterial cultures (PureLink Genomic DNA Mini Kit, Thermo Fisher) or stool (QIAamp Fast DNA Stool Mini Kit, Qiagen). Primers amplifying the VR region of each *Bacteroides* strain were designed with the forward primer containing a 20 bp random Unique Molecular Index (UMI) and an adapter, as described previously^111^. Briefly, each forward primer was designed to anneal to the VR region at 62-64°C and was present in 1/10 the normal concentration. A second forward primer would anneal to the adapter of the first forward primer at a temperature of 68-70°C. The second forward primer and reverse primer contained partial Illumina adapter sequences. Cycling parameters included 1 cycle of 98°C for 3 mins, 2 cycles of 98°C for 15 secs, 62°C for 45 secs, and 72°C for 30 secs, followed by 38 cycles of 98°C for 15 secs, 70°C for 45 secs, followed by 72°C for 5 mins. PCR products were purified using SPRI Select beads (Beckman Coulter) and sent for EZ Amplicon-Seq (Azenta Life Sciences).

### Data processing of Amplicon-Seq reads

Raw reads were trimmed and contamination filtered with BBDuk using default parameters except a kmer length of 23 and mink length of 11. Data was then merged with BBMerge^112^ using default parameters. Merged reads were then aligned to VR by creating a custom blastn^99^ database created from the parental VR sequences of *Bfr, Bth, Bun, Bov,* and *Bfi* to allow for a large number of potential mismatches using the parameters for blastn word_size=8, reward=1, penalty=-1, evalue=1e-5, gapopen=6, gapextend=6, and perc_identity=50. UMI and VR sequences were then extracted from the aligned read and compiled together. UMIs that contained mismatching sequences were discarded. Reads were then analyzed for the number of adenine-mutations and non-adenine mutations compared to parental strains. Reads that had >50% of non-adenine mutations were also discarded. The number of divergent reads was calculated by summing the number of reads with greater than 1 (*i.e.* 2 or more) adenine-mutations compared to parental strain (to account for potential sequencing errors). When plotted on a log chart, a pseudocount of 1 mutated read was added to all samples.

### Total RNA extraction, RNA-Seq Library Preparation, and Illumina Sequencing

*Bacteroides* cultures were grown to mid-log phase (OD_600_ 0.3–0.5) and stationary phase (OD_600_ 0.7–1.0) in BHIS. Cells were flash-frozen in a dry ice-ethanol slurry, then pelleted by centrifugation at 16,000xg for 1 minute at 4°C. The bacterial pellets were stored at −80°C until RNA extraction. RNA was extracted using TRIzol reagent and the PureLink™ RNA Mini Kit following the manufacturer’s instructions. Briefly, bacterial pellets were incubated with 1 mL of TRIzol for 5 minutes at room temperature, followed by the addition of 200 µL of chloroform. After 5 minutes at room temperature, the samples were centrifuged at 12,000xg for 15 minutes at 4°C. The aqueous phase was transferred to a new tube, and an equal volume of 100% ethanol was added. This mixture was transferred to a spin cartridge and centrifuged at 12,000xg for 15 seconds at room temperature.

RNA samples were treated with PureLink™ DNase I, according to the manufacturer’s protocol. Following DNase treatment and four wash steps, RNA was eluted in 50 µL of RNase-free water by centrifugation at 12,000xg for 1 minute. Eluted RNA was transferred to RNase-free tubes and analyzed for quality by agarose gel electrophoresis. RNA integrity was assessed using the Bio-Rad ChemiDoc system, and RNA concentration was quantified with a NanoDrop One spectrophotometer (Thermo Fisher Scientific). Further RNA quality and concentration was validated on an Agilent 4200 TapeStation RNA ScreenTape, with all RIN scores above 8.0 and samples normalized to 250 ng starting mass. Library preparation was performed using Illumina’s Stranded Total RNA Prep with Ribo-Zero Plus Microbiome, which targets depletion of microbiome rRNA prior to cDNA synthesis and library formation. Final libraries were validated using an Agilent 4200 TapeStation D1000 ScreenTape, with an average fragment size of 396bp. Libraries were quantified using Invitrogen’s Quant iT High Sensitivity dsDNA Kit and normalized during pooling. Samples were sequenced in 3 lanes of NovaSeq X Plus 10B PE 2x100.

### *Bacteroides* dataset and Mother-Infant dataset acquisition

The NCBI Assembly database was searched for the term “Bacteroides” and filtered to include those assemblies within the RefSeq database on 01/22/2021. Genome strain names were derived from the genome assembly reports. For the mother-infant dataset, raw reads were downloaded from BioProject PRJNA475246^85^ or metagenomic assemblies^84,96,97^ were downloaded as previously described^113^. Raw reads had adapters trimmed and were quality filtered using BBDuk and merged using BBMerge^112^. Merged reads were assembled into contigs using MegaHit^29^ using parameters: --min-contig-len 500 -m 0.85 --presets meta-sensitive. Contigs with length <2000 bp were then discarded and the resulting contigs were used for DGR identification. Taxonomic classification of contigs was performed by mmseqs2 using default parameters^114^.

### DGR identification from genomic and metagenomic datasets

A profile of previously identified DGR RT proteins^12,115^ was built using HMMER^56^. Predicted proteomes were derived from genomes and contigs using Prodigal^98^ and the resulting predicted proteins were searched for hits using HMMscan. A 20 kb window on either side of RT hits was used as input to search for imperfect repeats using a custom BLAST script^99^. Pairs were checked for one pair to be contained within a previously predicted ORF. Because there are many potential mismatches of reads to the VR sequence, traditional local aligners, such as Bowtie^116^ or BWA^117^ will trim or discard the read. To circumvent this challenge and determine DGR activity from metagenomic data, raw reads were mapped back to identified VRs by creating a custom BLAST database of the VR sequence using the same blastn parameters as above. Reads that fully or partially aligned were then checked for alignment to TR and the rest of the genome/contig by identifying if those reads had fewer mismatches than to VR. Any read which best aligned to the VR region was kept. Reads aligning to VR regions were then analyzed to determine mutations existed that corresponded to TR adenine positions. If adenine-specific mutations were found, the DGR was categorized as active. VR haplotypes were generated for each timepoint by generating the consensus VR sequence. The consensus sequence was compared between timepoints to determine if it differed.

### Identification of non-DGR RTs

A previously built profile of all RTs in bacterial genomes^51^ was modified to exclude the DGR RTs and rebuilt using HMMER. Proteomes from genomes or metagenomic contigs were searched for these RTs. The number of non-DGR RTs was reduced by random sampling and this subset was added to the identified DGR RTs for phylogenetic analysis.

### Clustering of variable proteins and phylogenetic tree building

In order to cluster variable proteins, each primary amino acid sequence was used for an all-vs-all blastp^99^ search. The result of the blastp was used as an edge weight for input into MCL clustering with inflation value of 2.0^101^. Within each cluster, a multiple sequence alignment was performed using MUSCLE^118^. This alignment was input into HHpred^54^ to search for functional domains within the proteins using the databases PDB_mmCIF70_17_Apr, Pfam-A_v35, and COG-KOG_v1.0. The RT phylogenetic tree was created by aligning each RT protein sequence, which was input into Fasttree2 using default parameters^100^. Variable proteins were classified as pilus proteins if they shared a significant domain with the following PBDs: 4EPS^48^, 4QB7^48^, 6JZJ^63^, 5NF4^119^. Variable proteins were classified as TaqVP-like if they shared a significant domain with PDB: 5VF4^57^. Variable proteins were classified as phage receptor tail binding proteins if they shared a significant domain with PDB: 1YU0^15^.

### DGR genomic assignments

Contigs containing DGRs were used as input for geNomad^52^, which predicts regions of contigs to be viral or plasmid. Coordinates from those predictions were aligned to DGR coordinates. DGRs located on a contig predicted to be entirely viral were classified as virus. DGRs located within a viral region surrounded by bacterial genes were classified as prophage. A 100 kb window surrounding DGRs located within predicted plasmid regions was then used as input for ICEfinder^53^ for ICE prediction. ICE synteny figures were generated using pyGenomeViz^120^.

### DGR gene transcription analysis

Raw sequencing reads from RNA-seq experiments were quality filtered and trimmed with fastp v0.23.4^121^ with default parameters, then mapped to the reference genome of each respective species with bwa-mem2^122^ with default parameters. Mapping statistics were generated with bamtools^123^ v2.5.1 and visualized with multiqc^124^ v1.21. Reads in gene features were counted with featureCounts^125^ function of the subread package at the fragment level (--countReadPairs) with fragments overlapping multiple features counted in both (-O). Total transcript counts were normalized to trimmed mean of M (TMM) values using NOISeq^126^ with default parameters. Within each *Bacteroides* strain, DGR gene TMM values were normalized to *gyrA* gene TMM values. The ratio of DGR genes to *gyrA* were then compared across strains, setting *Bfr* ratios to 1 to generate relative ratios for each of the other strains.

### VR mutagenesis and entropy analysis

VR mutagenesis frequency was calculated by taking the number of reads with >1 substitution from the canonical sequence divided by the total number of sequences. VR entropy was calculated using the formula:

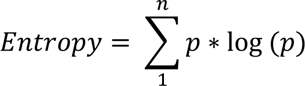

according to Shannon^81^, where p is the frequency of a unique VR sequence and n is the total number of unique mutagenized VR sequences.

### Statistical tests and metrics

Statistical comparisons were performed using the tests indicated in the figure legends. P-values were generated using ANOVA or student’s t-test. Multiple comparison corrections were performed using Holm-Sidak method or Tukey’s method, where appropriate. Statistical significance was defined at α=0.05.

### Data and code availability

Python code to identify DGRs from genomes will be made available at https://github.com/macadangdanglab/dgrdiscovery upon publication. Amplicon-Seq reads of VRs will be accessible on the short read archive (SRA) upon publication.

## Notes

### Competing Interest Statement

The authors have declared no competing interest.

## References

1. Heilbron, K., Toll-Riera, M., Kojadinovic, M., and MacLean, R.C. (2014). Fitness Is Strongly Influenced by Rare Mutations of Large Effect in a Microbial Mutation Accumulation Experiment. Genetics 197, 981–990. 10.1534/genetics.114.163147.

2. Mazel, D. (2006). Integrons: agents of bacterial evolution. Nat Rev Microbiol 4, 608–620. 10.1038/nrmicro1462.

3. Müller, F., and Tobler, H. (2000). Chromatin diminution in the parasitic nematodes Ascaris suum and Parascaris univalens. Int J Parasitol 30, 391–399. 10.1016/s0020-7519(99)00199-x.

4. Schatz, D.G., and Swanson, P.C. (2011). V(D)J Recombination: Mechanisms of Initiation. Annu Rev Genet 45, 167–202. 10.1146/annurev-genet-110410-132552.

5. Fitzgerald, D.M., and Rosenberg, S.M. (2019). What is mutation? A chapter in the series: How microbes “jeopardize” the modern synthesis. Plos Genet 15, e1007995. 10.1371/journal.pgen.1007995.

6. Shee, C., Gibson, J.L., Darrow, M.C., Gonzalez, C., and Rosenberg, S.M. (2011). Impact of a stress-inducible switch to mutagenic repair of DNA breaks on mutation in Escherichia coli. Proc National Acad Sci 108, 13659–13664. 10.1073/pnas.1104681108.

7. Jiang, F., and Doudna, J.A. (2015). CRISPR–Cas9 Structures and Mechanisms. Annu. Rev. Biophys. 46, 1–25. 10.1146/annurev-biophys-062215-010822.

8. Marraffini, L.A., and Sontheimer, E.J. (2008). CRISPR Interference Limits Horizontal Gene Transfer in Staphylococci by Targeting DNA. Science 322, 1843–1845. 10.1126/science.1165771.

9. Wheatley, R.M., and MacLean, R.C. (2021). CRISPR-Cas systems restrict horizontal gene transfer in Pseudomonas aeruginosa. ISME J. 15, 1420–1433. 10.1038/s41396-020-00860-3.

10. Macadangdang, B.R., Makanani, S.K., and Miller, J.F. (2022). Accelerated Evolution by Diversity-Generating Retroelements. Annu Rev Microbiol 76, 389–411. 10.1146/annurev-micro-030322-040423.

11. Doré, H., Eisenberg, A.R., Junkins, E.N., Leventhal, G.E., Ganesh, A., Cordero, O.X., Paul, B.G., Valentine, D.L., O’Malley, M.A., and Wilbanks, E.G. (2024). Targeted hypermutation of putative antigen sensors in multicellular bacteria. Proc. Natl. Acad. Sci. 121, e2316469121. 10.1073/pnas.2316469121.

12. Wu, L., Gingery, M., Abebe, M., Arambula, D., Czornyj, E., Handa, S., Khan, H., Liu, M., Pohlschroder, M., Shaw, K.L., et al. (2017). Diversity-generating retroelements: natural variation, classification and evolution inferred from a large-scale genomic survey. Nucleic Acids Res 46, gkx1150-. 10.1093/nar/gkx1150.

13. Miller, J.L., Coq, J.L., Hodes, A., Barbalat, R., Miller, J.F., and Ghosh, P. (2008). Selective Ligand Recognition by a Diversity-Generating Retroelement Variable Protein. Plos Biol 6, e131. 10.1371/journal.pbio.0060131.

14. Coq, J.L., and Ghosh, P. (2011). Conservation of the C-type lectin fold for massive sequence variation in a Treponema diversity-generating retroelement. Proc National Acad Sci 108, 14649–14653. 10.1073/pnas.1105613108.

15. McMahon, S.A., Miller, J.L., Lawton, J.A., Kerkow, D.E., Hodes, A., Marti-Renom, M.A., Doulatov, S., Narayanan, E., Sali, A., Miller, J.F., et al. (2005). The C-type lectin fold as an evolutionary solution for massive sequence variation. Nat Struct Mol Biol 12, 886–892. 10.1038/nsmb992.

16. Guo, H., Tse, L.V., Barbalat, R., Sivaamnuaiphorn, S., Xu, M., Doulatov, S., and Miller, J.F. (2008). Diversity-Generating Retroelement Homing Regenerates Target Sequences for Repeated Rounds of Codon Rewriting and Protein Diversification. Mol Cell 31, 813–823. 10.1016/j.molcel.2008.07.022.

17. Doulatov, S., Hodes, A., Dai, L., Mandhana, N., Liu, M., Deora, R., Simons, R.W., Zimmerly, S., and Miller, J.F. (2004). Tropism switching in Bordetella bacteriophage defines a family of diversity-generating retroelements. Nature 431, 476–481. 10.1038/nature02833.

18. Liu, M., Deora, R., Doulatov, S.R., Gingery, M., Eiserling, F.A., Preston, A., Maskell, D.J., Simons, R.W., Cotter, P.A., Parkhill, J., et al. (2002). Reverse Transcriptase-Mediated Tropism Switching in Bordetella Bacteriophage. Science 295, 2091–2094. 10.1126/science.1067467.

19. Roux, S., Paul, B.G., Bagby, S.C., Nayfach, S., Allen, M.A., Attwood, G., Cavicchioli, R., Chistoserdova, L., Gruninger, R.J., Hallam, S.J., et al. (2021). Ecology and molecular targets of hypermutation in the global microbiome. Nat Commun 12, 3076. 10.1038/s41467-021-23402-7.

20. Macadangdang, B.R., Makanani, S.K., and Miller, J.F. (2022). Accelerated Evolution by Diversity-Generating Retroelements. Annu Rev Microbiol 76. 10.1146/annurev-micro-030322-040423.

21. Faith, J.J., Guruge, J.L., Charbonneau, M., Subramanian, S., Seedorf, H., Goodman, A.L., Clemente, J.C., Knight, R., Heath, A.C., Leibel, R.L., et al. (2013). The Long-Term Stability of the Human Gut Microbiota. Science 341, 1237439. 10.1126/science.1237439.

22. Charbonneau, M.R., O’Donnell, D., Blanton, L.V., Totten, S.M., Davis, J.C.C., Barratt, M.J., Cheng, J., Guruge, J., Talcott, M., Bain, J.R., et al. (2016). Sialylated Milk Oligosaccharides Promote Microbiota-Dependent Growth in Models of Infant Undernutrition. Cell 164, 859–871. 10.1016/j.cell.2016.01.024.

23. Mazmanian, S.K., Round, J.L., and Kasper, D.L. (2008). A microbial symbiosis factor prevents intestinal inflammatory disease. Nature 453, 620–625. 10.1038/nature07008.

24. Portincasa, P., Bonfrate, L., Vacca, M., Angelis, M.D., Farella, I., Lanza, E., Khalil, M., Wang, D.Q.-H., Sperandio, M., and Ciaula, A.D. (2022). Gut Microbiota and Short Chain Fatty Acids: Implications in Glucose Homeostasis. Int. J. Mol. Sci. 23, 1105. 10.3390/ijms23031105.

25. Jean, S., Wallace, M.J., Dantas, G., and Burnham, C.-A.D. (2022). Time for Some Group Therapy: Update on Identification, Antimicrobial Resistance, Taxonomy, and Clinical Significance of the Bacteroides fragilis Group. J. Clin. Microbiol. 60, e02361–20. 10.1128/jcm.02361-20.

26. Schloissnig, S., Arumugam, M., Sunagawa, S., Mitreva, M., Tap, J., Zhu, A., Waller, A., Mende, D.R., Kultima, J.R., Martin, J., et al. (2013). Genomic variation landscape of the human gut microbiome. Nature 493, 45–50. 10.1038/nature11711.

27. Liebert, C.A., Hall, R.M., and Summers, A.O. (1999). Transposon Tn 21, Flagship of the Floating Genome. Microbiol Mol Biol R 63, 507–522. 10.1128/mmbr.63.3.507-522.1999.

28. Frost, L.S., Leplae, R., Summers, A.O., and Toussaint, A. (2005). Mobile genetic elements: the agents of open source evolution. Nat Rev Microbiol 3, 722–732. 10.1038/nrmicro1235.

29. Li, D., Liu, C.-M., Luo, R., Sadakane, K., and Lam, T.-W. (2015). MEGAHIT: an ultra-fast single-node solution for large and complex metagenomics assembly via succinct de Bruijn graph. Bioinformatics 31, 1674–1676. 10.1093/bioinformatics/btv033.

30. Bankevich, A., Nurk, S., Antipov, D., Gurevich, A.A., Dvorkin, M., Kulikov, A.S., Lesin, V.M., Nikolenko, S.I., Pham, S., Prjibelski, A.D., et al. (2012). SPAdes: A New Genome Assembly Algorithm and Its Applications to Single-Cell Sequencing. J Comput Biol 19, 455–477. 10.1089/cmb.2012.0021.

31. Nurk, S., Meleshko, D., Korobeynikov, A., and Pevzner, P.A. (2017). metaSPAdes: a new versatile metagenomic assembler. Genome Res 27, 824–834. 10.1101/gr.213959.116.

32. Boisvert, S., Raymond, F., Godzaridis, É., Laviolette, F., and Corbeil, J. (2012). Ray Meta: scalable de novo metagenome assembly and profiling. Genome Biol 13, R122. 10.1186/gb-2012-13-12-r122.

33. Afiahayati, Sato, K., and Sakakibara, Y. (2015). MetaVelvet-SL: an extension of the Velvet assembler to a de novo metagenomic assembler utilizing supervised learning. Dna Res 22, 69–77. 10.1093/dnares/dsu041.

34. Namiki, T., Hachiya, T., Tanaka, H., and Sakakibara, Y. (2012). MetaVelvet: an extension of Velvet assembler to de novo metagenome assembly from short sequence reads. Nucleic Acids Res 40, e155– e155. 10.1093/nar/gks678.

35. Peng, Y., Leung, H.C.M., Yiu, S.M., and Chin, F.Y.L. (2012). IDBA-UD: a de novo assembler for single-cell and metagenomic sequencing data with highly uneven depth. Bioinformatics 28, 1420–1428. 10.1093/bioinformatics/bts174.

36. Treangen, T.J., Koren, S., Sommer, D.D., Liu, B., Astrovskaya, I., Ondov, B., Darling, A.E., Phillippy, A.M., and Pop, M. (2013). MetAMOS: a modular and open source metagenomic assembly and analysis pipeline. Genome Biol 14, R2. 10.1186/gb-2013-14-1-r2.

37. Kultima, J.R., Coelho, L.P., Forslund, K., Huerta-Cepas, J., Li, S.S., Driessen, M., Voigt, A.Y., Zeller, G., Sunagawa, S., and Bork, P. (2016). MOCAT2: a metagenomic assembly, annotation and profiling framework. Bioinformatics 32, 2520–2523. 10.1093/bioinformatics/btw183.

38. Eren, A.M., Esen, Ö.C., Quince, C., Vineis, J.H., Morrison, H.G., Sogin, M.L., and Delmont, T.O. (2015). Anvi’o: an advanced analysis and visualization platform for ‘omics data. Peerj 3, e1319. 10.7717/peerj.1319.

39. Wu, Y.-W., Simmons, B.A., and Singer, S.W. (2016). MaxBin 2.0: an automated binning algorithm to recover genomes from multiple metagenomic datasets. Bioinformatics 32, 605–607. 10.1093/bioinformatics/btv638.

40. Alneberg, J., Bjarnason, B.S., Bruijn, I. de, Schirmer, M., Quick, J., Ijaz, U.Z., Lahti, L., Loman, N.J., Andersson, A.F., and Quince, C. (2014). Binning metagenomic contigs by coverage and composition. Nat Methods 11, 1144–1146. 10.1038/nmeth.3103.

41. Lu, Y.Y., Chen, T., Fuhrman, J.A., and Sun, F. (2016). COCACOLA: binning metagenomic contigs using sequence COmposition, read CoverAge, CO-alignment and paired-end read LinkAge. Bioinformatics 33, btw290. 10.1093/bioinformatics/btw290.

42. Kang, D.D., Froula, J., Egan, R., and Wang, Z. (2015). MetaBAT, an efficient tool for accurately reconstructing single genomes from complex microbial communities. Peerj 3, e1165. 10.7717/peerj.1165.

43. Laczny, C.C., Sternal, T., Plugaru, V., Gawron, P., Atashpendar, A., Margossian, H.H., Coronado, S., Maaten, L. van der, Vlassis, N., and Wilmes, P. (2015). VizBin - an application for reference-independent visualization and human-augmented binning of metagenomic data. Microbiome 3, 1. 10.1186/s40168-014-0066-1.

44. Wu, Y.-W., and Ye, Y. (2011). A Novel Abundance-Based Algorithm for Binning Metagenomic Sequences Using l-tuples. J Comput Biol 18, 523–534. 10.1089/cmb.2010.0245.

45. Imelfort, M., Parks, D., Woodcroft, B.J., Dennis, P., Hugenholtz, P., and Tyson, G.W. (2014). GroopM: an automated tool for the recovery of population genomes from related metagenomes. Peerj 2, e603. 10.7717/peerj.603.

46. Wang, Y., Leung, H.C.M., Yiu, S.M., and Chin, F.Y.L. (2012). MetaCluster 5.0: a two-round binning approach for metagenomic data for low-abundance species in a noisy sample. Bioinformatics 28, i356– i362. 10.1093/bioinformatics/bts397.

47. Patil, K.R., Roune, L., and McHardy, A.C. (2012). The PhyloPythiaS Web Server for Taxonomic Assignment of Metagenome Sequences. Plos One 7, e38581. 10.1371/journal.pone.0038581.

48. Xu, Q., Shoji, M., Shibata, S., Naito, M., Sato, K., Elsliger, M.-A., Grant, J.C., Axelrod, H.L., Chiu, H.-J., Farr, C.L., et al. (2016). A Distinct Type of Pilus from the Human Microbiome. Cell 165, 690–703. 10.1016/j.cell.2016.03.016.

49. O’Leary, N.A., Wright, M.W., Brister, J.R., Ciufo, S., Haddad, D., McVeigh, R., Rajput, B., Robbertse, B., Smith-White, B., Ako-Adjei, D., et al. (2016). Reference sequence (RefSeq) database at NCBI: current status, taxonomic expansion, and functional annotation. Nucleic Acids Res 44, D733– D745. 10.1093/nar/gkv1189.

50. Vallota-Eastman, A., Arrington, E.C., Meeken, S., Roux, S., Dasari, K., Rosen, S., Miller, J.F., Valentine, D.L., and Paul, B.G. (2020). Role of diversity-generating retroelements for regulatory pathway tuning in cyanobacteria. Bmc Genomics 21, 664. 10.1186/s12864-020-07052-5.

51. Sharifi, F., and Ye, Y. (2021). Identification and classification of reverse transcriptases in bacterial genomes and metagenomes. Nucleic Acids Res 50, e29–e29. 10.1093/nar/gkab1207.

52. Camargo, A.P., Roux, S., Schulz, F., Babinski, M., Xu, Y., Hu, B., Chain, P.S.G., Nayfach, S., and Kyrpides, N.C. (2023). Identification of mobile genetic elements with geNomad. Nat. Biotechnol., 1–10. 10.1038/s41587-023-01953-y.

53. Liu, M., Li, X., Xie, Y., Bi, D., Sun, J., Li, J., Tai, C., Deng, Z., and Ou, H.-Y. (2019). ICEberg 2.0: an updated database of bacterial integrative and conjugative elements. Nucleic Acids Res 47, D660– D665. 10.1093/nar/gky1123.

54. Söding, J., Biegert, A., and Lupas, A.N. (2005). The HHpred interactive server for protein homology detection and structure prediction. Nucleic Acids Res 33, W244–W248. 10.1093/nar/gki408.

55. Paul, B.G., Bagby, S.C., Czornyj, E., Arambula, D., Handa, S., Sczyrba, A., Ghosh, P., Miller, J.F., and Valentine, D.L. (2015). Targeted diversity generation by intraterrestrial archaea and archaeal viruses. Nat Commun 6, 6585. 10.1038/ncomms7585.

56. Eddy, S.R. (2011). Accelerated Profile HMM Searches. Plos Comput Biol 7, e1002195. 10.1371/journal.pcbi.1002195.

57. Handa, S., Shaw, K.L., and Ghosh, P. (2019). Crystal structure of a Thermus aquaticus diversity-generating retroelement variable protein. Plos One 14, e0205618. 10.1371/journal.pone.0205618.

58. Nguyen, K.B., Sreelatha, A., Durrant, E.S., Lopez-Garrido, J., Muszewska, A., Dudkiewicz, M., Grynberg, M., Yee, S., Pogliano, K., Tomchick, D.R., et al. (2016). Phosphorylation of spore coat proteins by a family of atypical protein kinases. Proc National Acad Sci 113, E3482–E3491. 10.1073/pnas.1605917113.

59. Welch, J.L.M., Rossetti, B.J., Rieken, C.W., Dewhirst, F.E., and Borisy, G.G. (2016). Biogeography of a human oral microbiome at the micron scale. Proc National Acad Sci 113, E791–E800. 10.1073/pnas.1522149113.

60. Welch, J.L.M., Ramírez-Puebla, S.T., and Borisy, G.G. (2020). Oral Microbiome Geography: Micron-Scale Habitat and Niche. Cell Host Microbe 28, 160–168. 10.1016/j.chom.2020.07.009.

61. Alayyoubi, M., Guo, H., Dey, S., Golnazarian, T., Brooks, G.A., Rong, A., Miller, J.F., and Ghosh, P. (2013). Structure of the Essential Diversity-Generating Retroelement Protein bAvd and Its Functionally Important Interaction with Reverse Transcriptase. Structure 21, 266–276. 10.1016/j.str.2012.11.016.

62. Jumper, J., Evans, R., Pritzel, A., Green, T., Figurnov, M., Ronneberger, O., Tunyasuvunakool, K., Bates, R., Žídek, A., Potapenko, A., et al. (2021). Highly accurate protein structure prediction with AlphaFold. Nature 596, 583–589. 10.1038/s41586-021-03819-2.

63. Shibata, S., Shoji, M., Okada, K., Matsunami, H., Matthews, M.M., Imada, K., Nakayama, K., and Wolf, M. (2020). Structure of polymerized type V pilin reveals assembly mechanism involving protease-mediated strand exchange. Nat Microbiol 5, 830–837. 10.1038/s41564-020-0705-1.

64. Hospenthal, M.K., Costa, T.R.D., and Waksman, G. (2017). A comprehensive guide to pilus biogenesis in Gram-negative bacteria. Nat Rev Microbiol 15, 365–379. 10.1038/nrmicro.2017.40.

65. Götzke, H., Kilisch, M., Martínez-Carranza, M., Sograte-Idrissi, S., Rajavel, A., Schlichthaerle, T., Engels, N., Jungmann, R., Stenmark, P., Opazo, F., et al. (2019). The ALFA-tag is a highly versatile tool for nanobody-based bioscience applications. Nat Commun 10, 4403. 10.1038/s41467-019-12301-7.

66. Johnson, C.M., and Grossman, A.D. (2015). Integrative and Conjugative Elements (ICEs): What They Do and How They Work. Annu Rev Genet 49, 1–25. 10.1146/annurev-genet-112414-055018.

67. Durrant, M.G., Li, M.M., Siranosian, B.A., Montgomery, S.B., and Bhatt, A.S. (2020). A Bioinformatic Analysis of Integrative Mobile Genetic Elements Highlights Their Role in Bacterial Adaptation. Cell Host Microbe 27, 140–153.e9. 10.1016/j.chom.2019.10.022.

68. Franke, A.E., and Clewell, D.B. (1981). Evidence for a chromosome-borne resistance transposon (Tn916) in Streptococcus faecalis that is capable of “conjugal” transfer in the absence of a conjugative plasmid. J. Bacteriol. 145, 494–502. 10.1128/jb.145.1.494-502.1981.

69. Mays, T.D., Smith, C.J., Welch, R.A., Delfini, C., and Macrina, F.L. (1982). Novel antibiotic resistance transfer in Bacteroides. Antimicrob. Agents Chemother. 21, 110–118. 10.1128/aac.21.1.110.

70. Roberts, M.C., and Smith, A.L. (1980). Molecular characterization of “plasmid-free” antibiotic-resistant Haemophilus influenzae. J. Bacteriol. 144, 476–479. 10.1128/jb.144.1.476-479.1980.

71. Shoemaker, N.B., Smith, M.D., and Guild, W.R. (1980). DNase-resistant transfer of chromosomal cat and tet insertions by filter mating in pneumococcus. Plasmid 3, 80–87. 10.1016/s0147-619x(80)90036-0.

72. Botelho, J., and Schulenburg, H. (2021). The Role of Integrative and Conjugative Elements in Antibiotic Resistance Evolution. Trends Microbiol. 29, 8–18. 10.1016/j.tim.2020.05.011.

73. Jaworski, D.D., and Clewell, D.B. (1994). Evidence that coupling sequences play a frequency-determining role in conjugative transposition of Tn916 in Enterococcus faecalis. J. Bacteriol. 176, 3328–3335. 10.1128/jb.176.11.3328-3335.1994.

74. Park, J., and Salyers, A.A. (2011). Characterization of the Bacteroides CTnDOT Regulatory Protein RteC. J Bacteriol 193, 91–97. 10.1128/jb.01015-10.

75. Wexler, A.G., and Goodman, A.L. (2017). An insider’s perspective: Bacteroides as a window into the microbiome. Nat Microbiol 2, 17026. 10.1038/nmicrobiol.2017.26.

76. Russell, A.B., Wexler, A.G., Harding, B.N., Whitney, J.C., Bohn, A.J., Goo, Y.A., Tran, B.Q., Barry, N.A., Zheng, H., Peterson, S.B., et al. (2014). A Type VI Secretion-Related Pathway in Bacteroidetes Mediates Interbacterial Antagonism. Cell Host Microbe 16, 227–236. 10.1016/j.chom.2014.07.007.

77. Arambula, D., Wong, W., Medhekar, B.A., Guo, H., Gingery, M., Czornyj, E., Liu, M., Dey, S., Ghosh, P., and Miller, J.F. (2013). Surface display of a massively variable lipoprotein by a Legionella diversity-generating retroelement. Proc National Acad Sci 110, 8212–8217. 10.1073/pnas.1301366110.

78. Naorem, S.S., Han, J., Wang, S., Lee, W.R., Heng, X., Miller, J.F., and Guo, H. (2017). DGR mutagenic transposition occurs via hypermutagenic reverse transcription primed by nicked template RNA. Proc National Acad Sci 114, E10187–E10195. 10.1073/pnas.1715952114.

79. Handa, S., Reyna, A., Wiryaman, T., and Ghosh, P. (2020). Determinants of adenine-mutagenesis in diversity-generating retroelements. Nucleic Acids Res. 49, 1033–1045. 10.1093/nar/gkaa1240.

80. Handa, S., Jiang, Y., Tao, S., Foreman, R., Schinazi, R.F., Miller, J.F., and Ghosh, P. (2018). Template-assisted synthesis of adenine-mutagenized cDNA by a retroelement protein complex. Nucleic Acids Res 46, gky620-. 10.1093/nar/gky620.

81. Shannon, C.E. (1948). A mathematical theory of communication. Bell Syst Technical J 27, 379–423. 10.1002/j.1538-7305.1948.tb01338.x.

82. Konopiński, M.K. (2020). Shannon diversity index: a call to replace the original Shannon’s formula with unbiased estimator in the population genetics studies. PeerJ 8, e9391. 10.7717/peerj.9391.

83. Brand, M.W., Wannemuehler, M.J., Phillips, G.J., Proctor, A., Overstreet, A.-M., Jergens, A.E., Orcutt, R.P., and Fox, J.G. (2015). The Altered Schaedler Flora: Continued Applications of a Defined Murine Microbial Community. ILAR J. 56, 169–178. 10.1093/ilar/ilv012.

84. Ferretti, P., Pasolli, E., Tett, A., Asnicar, F., Gorfer, V., Fedi, S., Armanini, F., Truong, D.T., Manara, S., Zolfo, M., et al. (2018). Mother-to-Infant Microbial Transmission from Different Body Sites Shapes the Developing Infant Gut Microbiome. Cell Host Microbe 24, 133–145.e5. 10.1016/j.chom.2018.06.005.

85. Yassour, M., Jason, E., Hogstrom, L.J., Arthur, T.D., Tripathi, S., Siljander, H., Selvenius, J., Oikarinen, S., Hyöty, H., Virtanen, S.M., et al. (2018). Strain-Level Analysis of Mother-to-Child Bacterial Transmission during the First Few Months of Life. Cell Host Microbe 24, 146–154.e4. 10.1016/j.chom.2018.06.007.

86. Bäckhed, F., Ding, H., Wang, T., Hooper, L.V., Koh, G.Y., Nagy, A., Semenkovich, C.F., and Gordon, J.I. (2004). The gut microbiota as an environmental factor that regulates fat storage. Proc National Acad Sci 101, 15718–15723. 10.1073/pnas.0407076101.

87. Lloyd-Price, J., Mahurkar, A., Rahnavard, G., Crabtree, J., Orvis, J., Hall, A.B., Brady, A., Creasy, H.H., McCracken, C., Giglio, M.G., et al. (2017). Strains, functions and dynamics in the expanded Human Microbiome Project. Nature 550, 61–66. 10.1038/nature23889.

88. Zepeda-Rivera, M., Minot, S.S., Bouzek, H., Wu, H., Blanco-Míguez, A., Manghi, P., Jones, D.S., LaCourse, K.D., Wu, Y., McMahon, E.F., et al. (2024). A distinct Fusobacterium nucleatum clade dominates the colorectal cancer niche. Nature, 1–9. 10.1038/s41586-024-07182-w.

89. Yaffe, E., and Relman, D.A. (2020). Tracking microbial evolution in the human gut using Hi-C reveals extensive horizontal gene transfer, persistence and adaptation. Nat. Microbiol. 5, 343–353. 10.1038/s41564-019-0625-0.

90. Zahavi, L., Lavon, A., Reicher, L., Shoer, S., Godneva, A., Leviatan, S., Rein, M., Weissbrod, O., Weinberger, A., and Segal, E. (2023). Bacterial SNPs in the human gut microbiome associate with host BMI. Nat. Med. 29, 2785–2792. 10.1038/s41591-023-02599-8.

91. Zhao, S., Lieberman, T.D., Poyet, M., Kauffman, K.M., Gibbons, S.M., Groussin, M., Xavier, R.J., and Alm, E.J. (2019). Adaptive Evolution within Gut Microbiomes of Healthy People. Cell Host Microbe 25, 656–667.e8. 10.1016/j.chom.2019.03.007.

92. Garud, N.R., Good, B.H., Hallatschek, O., and Pollard, K.S. (2019). Evolutionary dynamics of bacteria in the gut microbiome within and across hosts. PLoS Biol. 17, e3000102. 10.1371/journal.pbio.3000102.

93. Wolff, R., and Garud, N.R. (2023). Pervasive selective sweeps across human gut microbiomes. bioRxiv, 2023.12.22.573162. 10.1101/2023.12.22.573162.

94. Valles-Colomer, M., Blanco-Míguez, A., Manghi, P., Asnicar, F., Dubois, L., Golzato, D., Armanini, F., Cumbo, F., Huang, K.D., Manara, S., et al. (2023). The person-to-person transmission landscape of the gut and oral microbiomes. Nature 614, 125–135. 10.1038/s41586-022-05620-1.

95. Liu, Q., Du, X., Hong, X., Li, T., Zheng, B., He, L., Wang, Y., Otto, M., and Li, M. (2015). Targeting Surface Protein SasX by Active and Passive Vaccination To Reduce Staphylococcus aureus Colonization and Infection. Infect Immun 83, 2168–2174. 10.1128/iai.02951-14.

96. Huttenhower, C., Gevers, D., Knight, R., Abubucker, S., Badger, J.H., Chinwalla, A.T., Creasy, H.H., Earl, A.M., FitzGerald, M.G., Fulton, R.S., et al. (2012). Structure, function and diversity of the healthy human microbiome. Nature 486, 207–214. 10.1038/nature11234.

97. Bäckhed, F., Roswall, J., Peng, Y., Feng, Q., Jia, H., Kovatcheva-Datchary, P., Li, Y., Xia, Y., Xie, H., Zhong, H., et al. (2015). Dynamics and Stabilization of the Human Gut Microbiome during the First Year of Life. Cell Host Microbe 17, 690–703. 10.1016/j.chom.2015.04.004.

98. Hyatt, D., Chen, G.-L., LoCascio, P.F., Land, M.L., Larimer, F.W., and Hauser, L.J. (2010). Prodigal: prokaryotic gene recognition and translation initiation site identification. Bmc Bioinformatics 11, 119. 10.1186/1471-2105-11-119.

99. Camacho, C., Coulouris, G., Avagyan, V., Ma, N., Papadopoulos, J., Bealer, K., and Madden, T.L. (2009). BLAST+: architecture and applications. Bmc Bioinformatics 10, 421. 10.1186/1471-2105-10-421.

100. Price, M.N., Dehal, P.S., and Arkin, A.P. (2010). FastTree 2 – Approximately Maximum-Likelihood Trees for Large Alignments. Plos One 5, e9490. 10.1371/journal.pone.0009490.

101. Enright, A.J., Dongen, S.V., and Ouzounis, C.A. (2002). An efficient algorithm for large-scale detection of protein families. Nucleic Acids Res 30, 1575–1584. 10.1093/nar/30.7.1575.

102. García-Bayona, L., and Comstock, L.E. (2019). Streamlined Genetic Manipulation of Diverse Bacteroides and Parabacteroides Isolates from the Human Gut Microbiota. Mbio 10, e01762–19. 10.1128/mbio.01762-19.

103. Ito, T., Gallegos, R., Matano, L.M., Butler, N.L., Hantman, N., Kaili, M., Coyne, M.J., Comstock, L.E., Malamy, M.H., and Barquera, B. (2020). Genetic and Biochemical Analysis of Anaerobic Respiration in Bacteroides fragilis and Its Importance In Vivo. Mbio 11, e03238–19. 10.1128/mbio.03238-19.

104. Lim, B., Zimmermann, M., Barry, N.A., and Goodman, A.L. (2017). Engineered Regulatory Systems Modulate Gene Expression of Human Commensals in the Gut. Cell 169, 547–558.e15. 10.1016/j.cell.2017.03.045.

105. Ahuja, U., Shokeen, B., Cheng, N., Cho, Y., Blum, C., Coppola, G., and Miller, J.F. (2016). Differential regulation of type III secretion and virulence genes in Bordetella pertussis and Bordetella bronchiseptica by a secreted anti-σ factor. Proc National Acad Sci 113, 2341–2348. 10.1073/pnas.1600320113.

106. Thein, M., Sauer, G., Paramasivam, N., Grin, I., and Linke, D. (2010). Efficient Subfractionation of Gram-Negative Bacteria for Proteomics Studies. J Proteome Res 9, 6135–6147. 10.1021/pr1002438.

107. Hughes, C.S., Moggridge, S., Müller, T., Sorensen, P.H., Morin, G.B., and Krijgsveld, J. (2019). Single-pot, solid-phase-enhanced sample preparation for proteomics experiments. Nat. Protoc. 14, 68–85. 10.1038/s41596-018-0082-x.

108. Cox, J., and Mann, M. (2008). MaxQuant enables high peptide identification rates, individualized p.p.b.-range mass accuracies and proteome-wide protein quantification. Nat Biotechnol 26, 1367–1372. 10.1038/nbt.1511.

109. Jimenez-Morales, D., Campos, A.R., Dollen, J.V., Krogan, N., and Swaney, D. artMS: Analytical R tools for Mass Spectrometry. https://bioconductor.org/packages/release/bioc/html/artMS.html.

110. Goedhart, J., and Luijsterburg, M.S. (2020). VolcaNoseR is a web app for creating, exploring, labeling and sharing volcano plots. Sci. Rep. 10, 20560. 10.1038/s41598-020-76603-3.

111. Fields, B., Moeskjær, S., Friman, V., Andersen, S.U., and Young, J.P.W. (2021). MAUI-seq: Metabarcoding using amplicons with unique molecular identifiers to improve error correction. Mol Ecol Resour 21, 703–720. 10.1111/1755-0998.13294.

112. Bushnell, B., Rood, J., and Singer, E. (2017). BBMerge – Accurate paired shotgun read merging via overlap. Plos One 12, e0185056. 10.1371/journal.pone.0185056.

113. Pasolli, E., Asnicar, F., Manara, S., Zolfo, M., Karcher, N., Armanini, F., Beghini, F., Manghi, P., Tett, A., Ghensi, P., et al. (2019). Extensive Unexplored Human Microbiome Diversity Revealed by Over 150,000 Genomes from Metagenomes Spanning Age, Geography, and Lifestyle. Cell 176, 649–662.e20. 10.1016/j.cell.2019.01.001.

114. Steinegger, M., and Söding, J. (2017). MMseqs2 enables sensitive protein sequence searching for the analysis of massive data sets. Nat. Biotechnol. 35, 1026–1028. 10.1038/nbt.3988.

115. Paul, B.G., Burstein, D., Castelle, C.J., Handa, S., Arambula, D., Czornyj, E., Thomas, B.C., Ghosh, P., Miller, J.F., Banfield, J.F., et al. (2017). Retroelement guided protein diversification abounds in vast lineages of bacteria and archaea. Nat Microbiol 2, 17045–17045. 10.1038/nmicrobiol.2017.45.

116. Langmead, B., and Salzberg, S.L. (2012). Fast gapped-read alignment with Bowtie 2. Nat Methods 9, 357–359. 10.1038/nmeth.1923.

117. Li, H., and Durbin, R. (2009). Fast and accurate short read alignment with Burrows–Wheeler transform. Bioinformatics 25, 1754–1760. 10.1093/bioinformatics/btp324.

118. Edgar, R.C. (2004). MUSCLE: a multiple sequence alignment method with reduced time and space complexity. Bmc Bioinformatics 5, 113. 10.1186/1471-2105-5-113.

119. Hall, M., Hasegawa, Y., Yoshimura, F., and Persson, K. (2018). Structural and functional characterization of shaft, anchor, and tip proteins of the Mfa1 fimbria from the periodontal pathogen Porphyromonas gingivalis. Sci. Rep. 8, 1793. 10.1038/s41598-018-20067-z.

120. Shimoyama, Y. pyGenomeViz: A genome visualization python package for comparative genomics. https://github.com/moshi4/pyGenomeViz.

121. Chen, S. (2023). Ultrafast one-pass FASTQ data preprocessing, quality control, and deduplication using fastp. iMeta 2, e107. 10.1002/imt2.107.

122. Md, V., Misra, S., Li, H., and Aluru, S. (2019). Efficient Architecture-Aware Acceleration of BWA-MEM for Multicore Systems. 2019 IEEE Int. Parallel Distrib. Process. Symp. (IPDPS) *00*, 314–324. 10.1109/ipdps.2019.00041.

123. Barnett, D.W., Garrison, E.K., Quinlan, A.R., Strömberg, M.P., and Marth, G.T. (2011). BamTools: a C++ API and toolkit for analyzing and managing BAM files. Bioinformatics 27, 1691–1692. 10.1093/bioinformatics/btr174.

124. Ewels, P., Magnusson, M., Lundin, S., and Käller, M. (2016). MultiQC: summarize analysis results for multiple tools and samples in a single report. Bioinformatics 32, 3047–3048. 10.1093/bioinformatics/btw354.

125. Liao, Y., Smyth, G.K., and Shi, W. (2014). featureCounts: an efficient general purpose program for assigning sequence reads to genomic features. Bioinformatics 30, 923–930. 10.1093/bioinformatics/btt656.

126. Tarazona, S., Furió-Tarí, P., Turrà, D., Pietro, A.D., Nueda, M.J., Ferrer, A., and Conesa, A. (2015). Data quality aware analysis of differential expression in RNA-seq with NOISeq R/Bioc package. Nucleic Acids Res. 43, e140–e140. 10.1093/nar/gkv711.

